# An Immunogold Single Extracellular Vesicular RNA and Protein (^Au^SERP) Biochip to Predict Responses to Immunotherapy in Non-Small Cell Lung Cancer Patients

**DOI:** 10.1101/2021.10.30.466609

**Authors:** Luong T. H. Nguyen, Xinyu Wang, Kwang Joo Kwak, Jingjing Zhang, Tamio Okimoto, Joseph Amann, Xilal Y. Rima, Min Jin Yoon, Takehito Shukuya, Nicole Walters, Yifan Ma, Donald Belcher, Hong Li, Andre F. Palmer, David P. Carbone, L. James Lee, Eduardo Reátegui

**Affiliations:** William G. Lowrie Department of Chemical and Biomolecular Engineering, The Ohio State University; Columbus OH 43210, USA; Spot Biosystems LLC; Palo Alto, CA, USA; Comprehensive Cancer Center, The Ohio State University; Columbus, OH 43210, USA; Department of Respiratory Medicine, Juntendo University; Tokyo, Japan

## Abstract

Conventional PD-L1 immunohistochemical tissue biopsies only predict 20~40% of non-small cell lung cancer (NSCLC) patients that will respond positively to anti-PD-1/PD-L1 immunotherapy. Herein, we present an immunogold biochip to quantify single extracellular vesicular RNA and protein (^Au^SERP) as a non-invasive alternative. With only 3 μL of serum, PD-1/PD-L1 proteins on the surface of extracellular vesicles (EVs) and EV PD-1/PD-L1 messenger RNA (mRNA) cargo were detected at a single-vesicle resolution and exceeded the sensitivities of ELISA and qRT-PCR by 1000 times. By testing a cohort of 27 non-responding and 27 responding NSCLC patients, ^Au^SERP indicated that the single-EV mRNA biomarkers surpass the single-EV protein biomarkers at predicting patient responses to immunotherapy. Dual single-EV PD-1/PD-L1 mRNA detection differentiated responders from non-responders with an accuracy of 72.2% and achieved an NSCLC diagnosis accuracy of 93.2%, suggesting the potential for ^Au^SERP to provide enhanced immunotherapy predictions and cancer diagnoses within the clinical setting.

## INTRODUCTION

The immune system responds to cancer *via* a complex network of cellular interactions in which cytotoxic T cells, helper T cells, and natural killer cells are activated and work in concert against tumor cells (*1*). However, many metastatic tumors have adopted methods to hijack immune checkpoints to evade immune recognition. The overexpression of programmed cell death ligand 1 (PD-L1) on the surface of tumor cells, which binds to programmed cell death protein 1 (PD-1) on T cells leading to a blockade of T cell activation, protects tumor cells from T cell-mediated killing (*2*). The manipulation of immune checkpoint pathways using immune checkpoint inhibitors (ICIs) has emerged as an effective form of immunotherapy, demonstrating positive and durable clinical outcomes (*3*). For instance, patients with metastatic melanoma treated with concurrent ipilimumab (anti-cytotoxic T lymphocyte-associated molecule-4 (anti-CTLA-4)) and nivolumab (anti-PD-1) achieved an overall survival rate of 79% in two years (*4*). However, a majority of cancer patients do not respond positively to immunotherapy. For example, the response rate to single-agent PD-1/PD-L1 inhibition in patients with renal cell carcinoma is only 19% (*5*). Hence, there is an urgency to determine which individual patients may benefit from PD-1/PD-L1 blockade and other immunotherapies.

Tumor PD-L1 expression has been approved by the Food and Drug Administration (FDA) as a predictive biomarker for anti-PD-1/PD-L1 immunotherapy, which is detected using immunohistochemistry (IHC) (*6*). Four PD-L1 IHC assays using four different anti-PD-L1 antibodies (22C3, 28–8, SP263, and SP142) on two different automated staining platforms (Dako and Ventana) have been registered with the FDA (*7, 8*). Patients with higher expression of PD-L1 on their tissue biopsies are associated with improved response rates to PD-1/PD-L1 blockade (*9*). However, sampling at a single metastatic site may not represent the entire tumor burden in a highly heterogeneous cancer (*10, 11*). Hence, it is desirable to develop novel technologies to detect predictive biomarkers from bodily fluids in a non-invasive manner. Such an approach can help integrate signals from all metastatic foci and can be repeated serially throughout immunotherapy with ease.

Circulating microRNAs (miRNAs) have been utilized as potential predictive biomarkers for anti-PD-1/PD-L1 immunotherapy for NSCLC (*12, 13*). However, precise quantification of circulating miRNAs is highly challenging due to inconsistencies originating from pre- and post-analytical variables and the inability to discriminate among closely related miRNAs (*14*). In this regard, circulating messenger RNAs (mRNAs), which can be protected within extracellular vesicles (EVs) from *in vivo* degradation, are superior to circulating miRNAs as practical predictive biomarkers for clinical use (*15*).

EVs are lipid particles released from cells that vary from 30 nm to a few microns in diameters and are present in all biological fluids (e.g., blood, urine, and cerebral spinal fluid) (*16*). EVs can be classified by size as “small EVs” (smaller than 200 nm) and “medium/large EVs” (larger than 200 nm), by biochemical composition, or by the environmental conditions of their biogenesis in accordance to the Minimal Information for Studies of Extracellular Vesicles (MISEV2018) guideline (*17*). EVs contain different cargo, including proteins, RNA, DNA, and lipids that can be trafficked between cells and serve as mediators of intercellular communication (*18*). There have been some efforts to characterize PD-1/PD-L1 proteins and PD-L1 mRNA from EVs using western blot (*19*), enzyme-linked immunosorbent assay (ELISA) (*20*), flow cytometry (*21*), quantitative reverse transcription polymerase chain reaction (qRT-PCR) (*22*), and digital droplet PCR (ddPCR) (*23*). However, these existing methods primarily focus on the bulk analysis of total proteins and RNA extracted from many EVs, lending averaged information, limiting the assay’s resolution and sensitivity (*24*). During the characterization process of bulk-analysis methods, EVs are broken down to obtain their internal contents, leading to the loss of molecular information at individual EV levels (*25*). Therefore, it is imperative to develop technologies that provide an accurate and efficient analysis of the molecular content at a single-EV level. Several single-EV analytical technologies have been proposed for EV molecular analysis, including analysis by fluorescence imaging of immobilized single EVs (*26–28*), flow cytometry of single EVs enlarged *via* target-initiated engineering (*29*), droplet-based single-exosome-counting enzyme-linked immunoassay (droplet digital ExoELISA) (*30*), digital detection integrated with surface-anchored nucleic acid amplification (*31*), proximity-dependent barcoding assay (*32*), and immuno-droplet digital polymerase chain reaction (iddPCR) (*33*). These methods successfully improved the limit of detection (LOD) and demonstrated heterogeneous protein profiles of single EVs unachievable by bulk-analysis methods. Despite this progress, detecting mRNAs and low abundance proteins (such as PD-1/PD-L1) in single EVs remains challenging due to the inherent limitations of signal-to-background thresholding (*34*).

Herein, we describe a novel technology that enables single-EV capture and detection to quantify low abundance biomarkers for immunotherapy. We engineered a highly sensitive immunogold-based biochip to co-quantify PD-1/PD-L1 proteins and mRNAs in single EVs sorted from non-small cell lung cancer (NSCLC) patient serum. Multiplexed analyses enabled the simultaneous detection of multiple targets in a single assay, thereby promoting the discovery of novel biomarker combinations for an improved diagnosis, prognosis, and prediction. Our gold nanoparticle-based single extracellular vesicular RNA and protein (^Au^SERP) biochip was formulated on a gold-coated glass coverslip functionalized with polyethylene glycol (PEG), to prevent non-specific binding, and gold spherical nanoparticles (NPs), to amplify the signal and improve single-EV sensitivity. Different antibodies were tethered on the chip surface to capture and sort EVs into subpopulations based on their membrane protein compositions. Anti-PD-1/PD-L1 antibodies and the tyramide signal amplification (TSA) method were used to quantify the corresponding membrane proteins on the captured single EVs. Molecular beacons (MBs) with target-specific probes were also fused with the captured single EVs to identify and quantify PD-1/PD-L1 mRNAs. Molecular characterization at the single-EV level was achieved by coupling ^Au^SERP that maximizes the signal-to-noise ratio (SNR) with high-resolution total internal reflection fluorescence (TIRF) microscopy. Serum samples from 27 NSCLC non-responders and 27 NSCLC responders collected before immunotherapy were analyzed for four single-EV biomarkers (PD-1/PD-L1 proteins and mRNAs). We showed that ^Au^SERP was highly sensitive to detecting single-EV protein and mRNA. Compared to single-EV protein biomarkers, single-EV mRNA cargo excelled at diagnosing NSCLC and predicting patient responses to immunotherapy. By combining single-EV PD-1/PD-L1 mRNA biomarkers, ^Au^SERP diagnosed NSCLC patients and predicted patient responses to anti-PD-1/PD-L1 immunotherapy with accuracies of 93.2% and 72.2%, respectively, suggesting the potential applications of ^Au^SERP as a non-invasive and competitive alternative to diagnose cancer patients and predict immunotherapy responses.

## RESULTS

### ^Au^SERP maximizes the signal-to-noise ratio and offers high-throughput analyses for single-EV detection

High-resolution TIRF microscopy has been widely used in cell biology to detect single molecules due to its high SNR and capability to detect single molecules (*35*). This imaging technique restricts excitation to a precise focal plane near the coverslip and eliminates out-of-focus fluorescence, allowing single-molecule detection (*36*). High-resolution TIRF microscopy has been applied to visualize EVs pre-stained with a fluorescent membrane dye on a glass slide (*35*). However, for protein-specific on-chip EV labeling, the non-specific binding of labeling antibodies to the glass coverslip complicates differentiating true binding events from the background noise. We designed a biochip with a PEG coating to prevent the non-specific binding of biomolecules to the glass surface. We first deposited a thin gold coating (thickness ~ 12 nm) onto a glass coverslip *via* titanium (thickness ~ 2 nm), which serves as a ‘metal glue’, and subsequently coated it with PEG (2 kDa). ^Au^SERP was then assembled by attaching the functionalized glass coverslip to a silicone gasket with 64 chambers (**Fig. 1a**). This design is highly scalable, allowing for up to four ^Au^SERP biochips to be placed into a tray for a high-throughput robotic processing of up to 256 samples. The treated glass surface was then functionalized with streptavidin-conjugated gold NPs and a cocktail of biotinylated antibodies to capture specific EV subpopulations (**Fig. 1b**). To further minimize non-specific binding, the treated glass was blocked with 3% (w/v) bovine serum albumin (BSA) and 0.05% (v/v) Tween-20 before and after EV capture. The thin gold coating was made to further improve the SNR of TIRF microscopy through the surface plasmon resonance (SPR) effect, which occurs when total internal reflection occurs at a metal film-liquid interface (*37, 38*). We previously showed that a biochip coated with a thin gold film and PEG could sensitively quantify target RNAs within EVs in bulk for an early non-invasive cancer diagnosis (*39*). In that design, EVs were captured on the biochip using cationic lipoplex nanoparticles (CLNs) *via* electrostatic fusion, and EV RNA cargos were detected with MBs encapsulated within the CLNs. However, the immobilization of highly positively charged CLNs (zeta potential ~ 30 mV) on the biochip surface caused significantly high background noise for RNA detection (*39*), which impairs the capability and sensitivity required for single-EV analyses. To enable both protein and RNA detections at a single-EV level, in the present study, we designed the novel ^Au^SERP biochip in which single EVs were captured and sorted *via* antibodies tethered onto gold NPs.

**Fig. 1.**
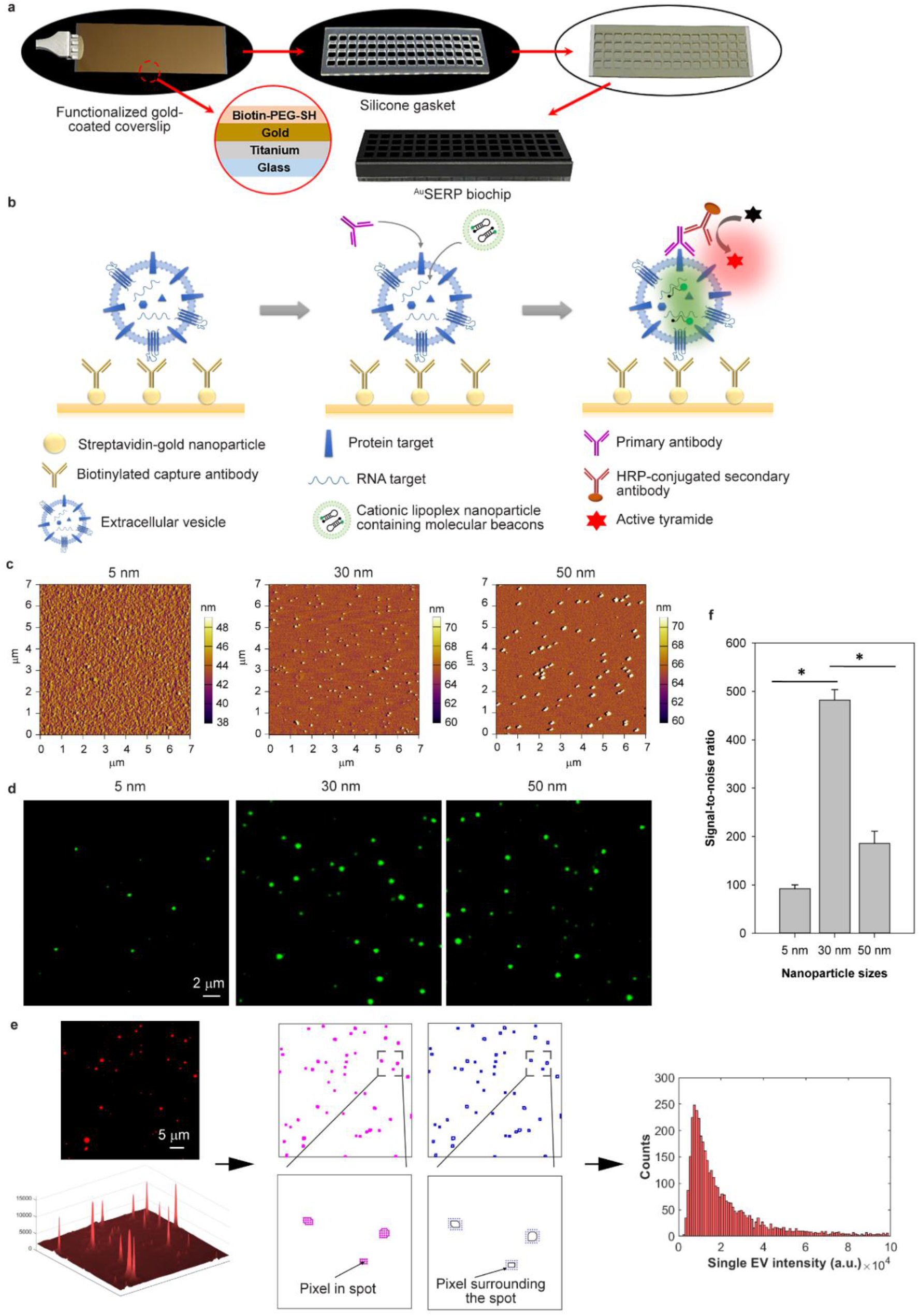
Design, characterization, and optimization of ^Au^SERP. **a** ^Au^SERP assembly. A functionalized gold-coated coverslip was attached to a silicone gasket with 64 chambers for high-throughput analysis of single-EV biomarkers. **b** A schematic mechanism for the detection of protein and mRNA biomarkers present in single extracellular vesicles (EVs) using ^Au^SERP. A gold-coated coverslip with PEG-tethered gold nanoparticles (NPs) conjugated to capture antibodies, was used to immobilize single EVs. Proteins on the surface of the single EVs were detected using the corresponding primary antibody and a tyramide signal amplification (TSA) method, resulting in fluorescent signals. mRNA cargo was identified using target-specific molecular beacons (MBs) encapsulated in cationic lipoplex nanoparticles (CLNs), resulting in fluorescent signals. **c** Atomic force microscopic (AFM) images of ^Au^SERP coated with different-sized gold NPs (5, 30, and 50 nm). **d** Representative total internal reflection fluorescence (TIRF) microscopic images of CD63 protein expression on the surface of H1568 single EVs captured with ^Au^SERP with different NP sizes. The images were cropped and enlarged from their original images, which are provided in Supplementary Fig. S1b. **e** Image processing workflow. Background noise signals surrounding bright spots were first removed using the Wavelet denoising method. All the pixels in each spot were then identified and subtracted by the mean intensity of pixels surrounding the spot. The noise-subtracted intensities of each pixel within the spot were summed into a net intensity of the single EV. Net intensities of all single EVs in 100 TIRF microscopic images were then collected to generate a histogram of net fluorescence intensity. **e** A signal-to-noise ratio (SNR) comparison of ^Au^SERP with different sized gold NPs for single-EV capture. The signal was calculated from the total fluorescence intensity of the CD63 surface protein levels on every single EV (individual fluorescence spot). The data were expressed as mean ± SD; n = 3; \**p* < 0.0001, Student’s *t*-test. a.u., arbitrary units.

We first investigated the effect of streptavidin-conjugated gold NP size (5, 30, and 50 nm) on the SNR of ^Au^SERP. Atomic force microscopic (AFM) images showed that NPs of all sizes were uniformly dispersed on surfaces (**Fig. 1c & Supplementary Fig. S1a**). The dispersion of NPs on ^Au^SERP promoted a single-EV resolution. With the same concentration loaded into each well (0.005% (w/v) based on gold), the number of NPs coated on the surface decreased as particle size increased. EVs from H1568 cells, an NSCLC cell line, were used to evaluate ^Au^SERP. The purified EVs were captured on the chip using a cocktail of anti-CD9 and anti-CD63 biotinylated antibodies. CD63 and CD9 are abundant tetraspanin surface antigens on EVs. CD63 staining of the captured EVs was used to evaluate the effect of NP sizes on the fluorescence signal intensity and background noise. TIRF images and histograms of net fluorescence intensity revealed that surfaces coated with 30-nm gold NPs had the highest signal and the lowest background, while both signal and background were lower for those coated with 5-nm NPs and higher for those coated with 50-nm NPs (**Fig. 1d & Supplementary Fig. S1b-c**). Fluorescence intensities of biomarker signals from TIRF images were quantified using a custom-built MATLAB algorithm, as shown in **Fig. 1e**, which allows for a rapid and automated single-EV image analysis. Briefly, our algorithm recognized a collective of spots with distinguished edges and removed background noise signals surrounding the bright spots using the Wavelet denoising method. All collectives of spots falling within a threshold of 3 – 8 pixels, set through a user interface, were identified as biomarker signals from single EVs (all other signals were discarded as EV aggregates or noise). All the pixels within each spot were identified and subtracted by the mean intensity of pixels surrounding the spot. The noise-subtracted intensities of each pixel within the spot were then summed into a net intensity of the single EV. Net intensities of all single EVs in 100 TIRF images were then collected to generate a histogram of net fluorescence intensity and calculate the total fluorescence intensity (TFI). Consequently, the SNR of ^Au^SERP was defined as

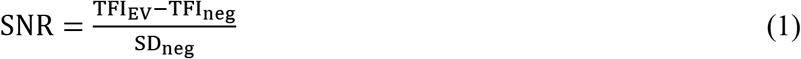

where TFI_EV_ is the TFI of signals obtained from EV samples, TFI_neg_ is the mean TFI of signals obtained from blanks (negative control – phosphate-buffered saline (PBS)), and SD_neg_ (or noise) is the standard deviation of signals obtained from blanks (by three independent experiments). Among the three sizes, 30 nm was shown to produce the highest SNR (*p* < 0.0001, **Fig. 1f**). Therefore, 30-nm gold NPs were chosen for our subsequent experiments. Our results were consistent with the observation that the size of gold NPs significantly affects the level of signal enhancement due to changes in surface coverage and refractive index (RI), where larger NPs resulted in greater RI shift (*40*).

### ^Au^SERP enables single-EV PD-L1 protein detection with ~ 1000 times more sensitivity than conventional ELISA

We next evaluated ^Au^SERP coated with 30-nm gold NPs for PD-1/PD-L1 protein and mRNA characterization of single EVs derived from *in vitro* cell model systems. The schematic of single-EV PD-1/PD-L1 protein and mRNA detection on ^Au^SERP is shown in **Fig. 1b**. A TSA method was employed to boost PD-1/PD-L1 protein signals on single-EV surfaces. This method uses the enzyme horseradish peroxidase (HRP) – a secondary antibody – to convert labeled tyramide molecules at the epitope detection site into a highly reactive oxidized intermediate, which binds rapidly and covalently to electron-rich tyrosine residues present in proteins near the epitope (*41*). Therefore, the TSA method can generate high-density labels of a target protein, making it 10 – 200 times more sensitive than conventional immunostaining methods. PD-1/PD-L1 mRNAs inside the single EVs were detected using CLNs encapsulating molecular beacons (MBs) (CLN-MBs) to provide a strong signal of target RNAs due to increased delivery efficiency of MBs into EVs compared to free MBs. As positively charged lipid nanoparticles, CLNs fuse with negatively charged EVs and deliver many MBs into the EVs (*42*). In contrast, the delivery of free MBs into EV lipid bilayers without any assistance is restricted to Brownian motion. Thus, our technology allows for the co-quantification of PD-1/PD-L1 molecular contents inside and on the surface of single EVs, thereby providing more comprehensive information of single EVs from NSCLC patients.

An *in vitro* PD-L1 model was developed by stimulating H1568 cells with interferon-gamma (IFN-γ), a cytokine secreted by activated effector T cells. IFN-γ is critical for innate and adaptive immunity and is known to upregulate PD-L1 expression on tumor cells (*43, 44*). In the present study, IFN-γ significantly increased the PD-L1 protein expression on H1568 cells as shown by IHC (**Fig. 2a**) and ELISA (*p* < 0.001, **Fig. 2b**). IHC was performed using an FDA-approved diagnostic assay, Dako PD-L1 IHC 22C3 pharmDx (*8*). An anti-PD-L1 antibody (Cell Signaling Technology (CST)) was also used to conduct PD-L1 immunofluorescence on the cells (**Supplementary Fig. S2a**). The result was comparable to the PD-L1 clinical assay, indicating the CST anti-PD-L1 antibody’s efficiency in binding to PD-L1 antigens. This antibody was, therefore, selected for further PD-L1 protein characterization on single EVs with ^Au^SERP.

**Fig. 2.**
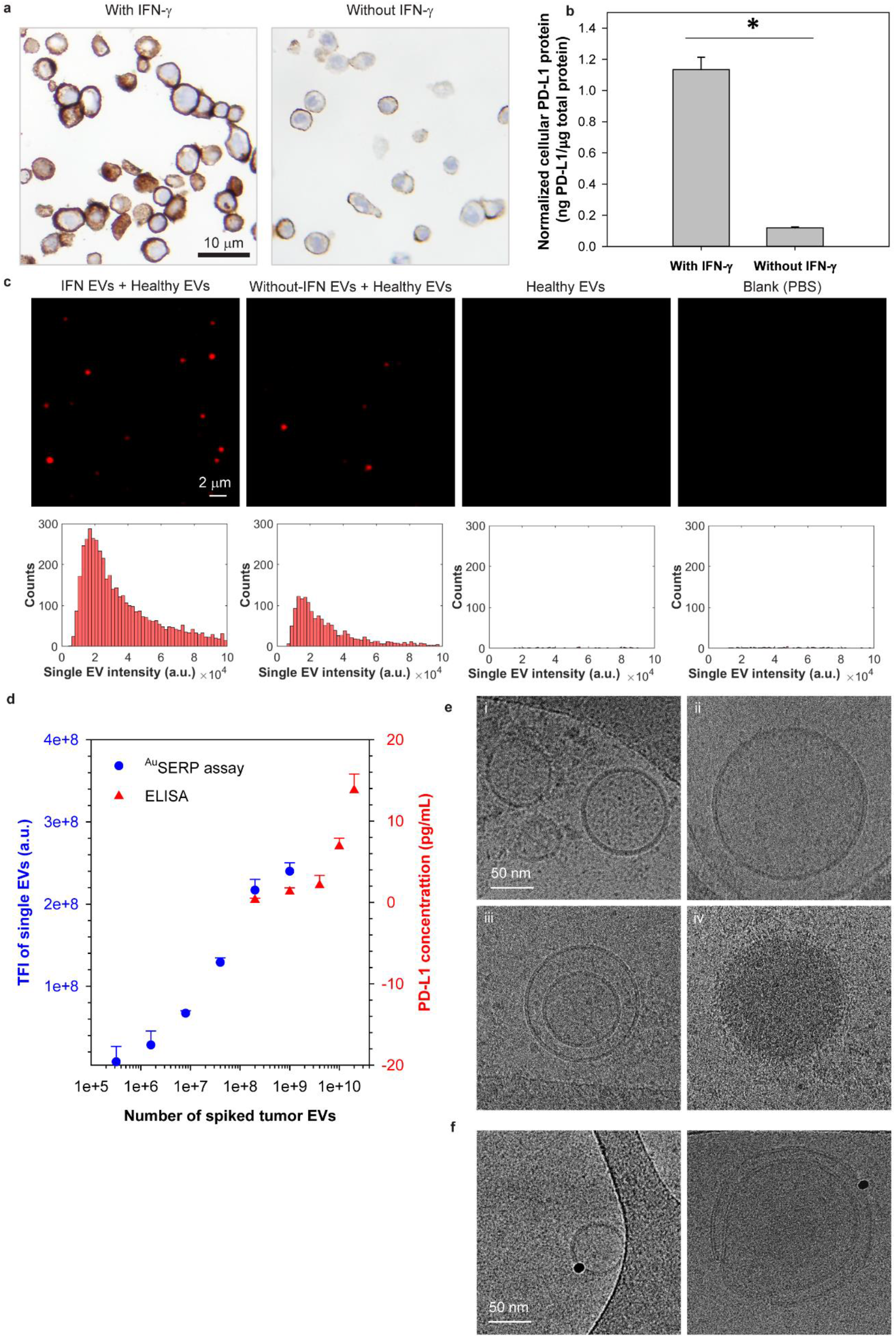
*In vitro* model and characterization of cellular and single-EV PD-L1 protein. **a** Immunohistochemistry (IHC) of PD-L1 protein in H1568 cells with/without interferon-gamma (IFN-γ) stimulation. Cell nuclei and PD-L1 protein were stained blue (by hematoxylin) and brown (by Dako PD-L1 IHC 22C3 pharmDx), respectively. **b** Quantification of PD-L1 protein levels in H1568 cells measured by ELISA and normalized by the total protein expressed by the cells measured with a BCA Protein Assay kit. The data were expressed as mean ± SD; n = 3; \**p* < 0.001, Student’s *t*-test. **c** Representative TIRF microscopic images and their corresponding histograms of PD-L1 protein expression on the surface of single EVs derived from H1568 cells with/without IFN-γ stimulation in comparison to healthy donor EVs and PBS as controls. The single-EV PD-L1 protein signals were characterized with ^Au^SERP using anti-PD-L1 antibodies and the TSA method. H1568 EVs were spiked in healthy donor EVs at a 1:1 ratio with 5 × 10^10^ particles/mL each. Healthy donor EVs were purified from healthy donor serum and then diluted in PBS to reach the target concentration. The images were cropped and enlarged from their original images, which are provided in Supplementary Fig. S2b. **d** A performance evaluation of the ^Au^SERP for PD-L1 protein detection in comparison to ELISA. EVs derived from IFN-γ-stimulated H1568 cells were spiked in healthy donor EVs at different concentrations ranging from 0 to 5 × 10^10^ particles/mL. The healthy donor EV concentration was kept constant at 5 × 10^10^ EVs/mL for all samples. The limit of detection (LOD) of ^Au^SERP for PD-L1 protein was ~ 10^6^ spiked tumor EVs, ~ 1000 times lower than ELISA. The data were expressed as mean ± SD; n = 3. TFI, total fluorescence intensity; a.u., arbitrary units. **e** Cryo-TEM images of EVs produced by IFN-γ-stimulated H1568 cells. **f** Cryo-TEM images of immunogold labeled PD-L1 protein on the EV surface.

PD-L1 protein on the surface of single EVs produced by H1568 cells with/without IFN-γ stimulation was characterized using ^Au^SERP with the CST anti-PD-L1 antibody. H1568 EVs were spiked in healthy donor EVs at a 1:1 ratio (H1568 EVs: healthy donor EVs) with 5 × 10^10^ particles/mL each to characterize PD-L1 expression. Healthy donor EVs at 5 × 10^10^ particles/mL and PBS blank samples were also examined as negative controls. EV staining procedures were performed without a permeabilization buffer to preserve the PD-L1 protein on the single-EV membrane surface. TIRF images and their corresponding histograms showed that PD-L1 protein expression on H1568 EVs with/without IFN-γ stimulation were successfully detected with ^Au^SERP and were significantly greater than the negative controls (*p* < 0.001, **Fig. 2c & Supplementary Fig. S2b**). The minimal signal detected in the blank control (PBS) may be derived from the substrate autofluorescence and/or non-specific binding of detection antibodies, which were insignificant compared to the true signal.

Additionally, the PD-L1 protein expression on single EVs from IFN-γ-stimulated H1568 cells was considerably higher than single EVs derived from cells without stimulation (*p* < 0.005, **Fig. 2c & Supplementary Fig. S2b**). Similar to cells, previous studies revealed that the levels of PD-L1 protein on tumor-derived EVs were also upregulated following IFN-γ stimulation (*19, 20*). Thus, our platform was sensitive enough to differentiate PD-L1 protein levels on single EVs of these different conditions. To indicate the robustness of ^Au^SERP for single-EV characterization, we tested an Abcam anti-PD-L1 antibody (clone 28-8) and compared it with the CST anti-PD-L1 antibody. Total fluorescence intensities of PD-L1 protein signals provided by both antibodies were comparable (*p* > 0.05, **Supplementary Fig. S2c**). However, there was a slight qualitative difference, where larger spot sizes and fewer spots were detected by the Abcam antibody (**Supplementary Fig. S2d**).

We then compared the LOD of ^Au^SERP with the most sensitive commercial ELISA kit, which has a LOD of 0.6 pg/mL. EVs produced from IFN-γ-stimulated H1568 cells were spiked into healthy donor EVs at different concentrations ranging from 0 to 5 × 10^10^ particles/mL and quantified for PD-L1 protein expression using ^Au^SERP and ELISA. The healthy donor EV concentration was kept constant at 5 × 10^10^ EVs/mL for all samples. While the minimum sensitivity of ELISA was ~ 10^9^ spiked H1568 EVs, our system could detect as low as ~ 10^6^ spiked H1568 EVs (**Fig. 2d**). As such, our platform outperformed ELISA in analytical sensitivity by ~ 1000 times. The LOD of our system was determined using the SNR method, in which an SNR of three is generally accepted for estimating the LOD (*45, 46*).

We also characterized the size, concentration, morphology, and structure of H1568 EVs to understand their physical properties. Size distributions of EVs with/without IFN-γ stimulation measured by tunable resistive pulse sensing (TRPS) are shown in **Supplementary Fig. S3a-b**. In both cases, most EVs were ‘small EVs’. There was also no significant difference in the number of EVs produced by the cells with/without IFN-γ stimulation (*p* > 0.05, **Supplementary Fig. S3c**). As such, IFN-γ did not increase the number of EVs secreted by the cells, consistent with a previous report (*47*), but upregulated the level of PD-L1 protein on EVs (**Fig. 2c & Supplementary Fig. S2b**). EVs isolated from H1568 cells with IFN-γ stimulation were also observed using the cryogenic transmission electron microscopy (cryo-TEM) technique. Most of the EVs were intact and had a round shape with a clear lipid bilayer/membrane and a translucent internal structure, indicating little material within the lumen (**Fig. 2e/i-ii & Supplementary Fig. S3d/i**). However, some particles had a round shape with electron-dense cargo and no visible lipid membrane (**Fig. 2e/iv**). Single, double, and multilayer vesicles with different sizes were also visualized (**Fig. 2e/i-iii & Supplementary Fig. S3d**). These observations demonstrate the vast heterogeneity of EVs, as mentioned in previous studies (*48, 49*). The presence of PD-L1 protein on the surface membrane of single EVs was also confirmed by immunogold labeling with cryo-TEM imaging, where single gold NPs bound to the membranes of single EVs *via* PD-L1 (**Fig. 2f**).

### ^Au^SERP enables single-EV PD-L1 mRNA detection with ~ 1000 times more sensitivity than conventional qRT-PCR

The presence of PD-L1 mRNA in plasma-derived EVs of NSCLC patients was previously demonstrated using ddPCR (*23*). Therefore, in addition to protein characterization, we aimed at detecting mRNAs inside the single EVs with ^Au^SERP. We first designed a PD-L1 MB and examined its fluorescent *in situ* hybridization capabilities with PD-L1 mRNA in H1568 cells. The PD-L1 MBs successfully localized PD-L1 mRNAs in the cytoplasm, with stronger signals in the IFN-γ-stimulated cells compared to the cells without IFN-γ stimulation (**Fig. 3a**). This result was consistent with PD-L1 mRNA quantification by qRT-PCR in which cellular PD-L1 mRNA was significantly increased with IFN-γ stimulation (*p* < 0.0001, **Fig. 3b**).

**Fig. 3.**
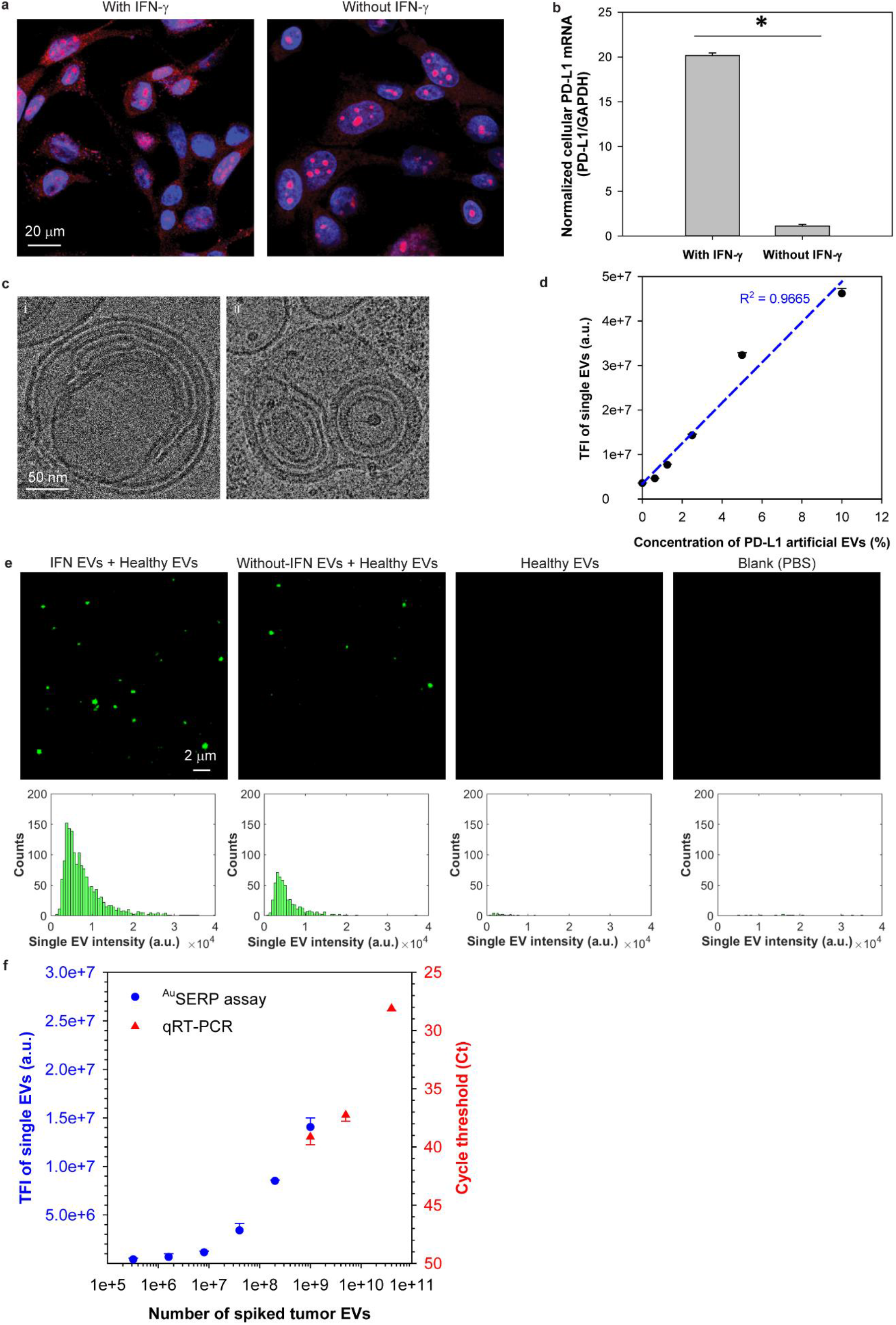
*In vitro* model and characterization of cellular and single-EV PD-L1 mRNA. **a** Confocal fluorescence microscopic images of PD-L1 mRNA in H1568 cells with/without IFN-γ stimulation. Cell nuclei and PD-L1 mRNA were stained blue (by DAPI) and red (by PD-L1 MBs via fluorescent *in situ* hybridization), respectively. **b** qRT-PCR analysis of PD-L1 mRNA levels in H1568 cells with/without IFN-γ stimulation. The data were expressed as mean ± SD; n = 3; \**p* < 0.0001, Student’s *t*-test. **c** Cryo-TEM images of a CLN-MB with a typical ‘onion-like structure (**i**) and the fusion of EVs produced by IFN-γ-stimulated H1568 cells with CLN-MBs targeting PD-L1 (**ii**). **d** Calibration of PD-L1-targeting CLN-MBs using artificial EVs made of liposomes encapsulating PD-L1 ssDNA oligos. There is a strong linear relationship between the resulting fluorescence signal and the concentration of artificial EVs (R^2^ = 0.9665, *p* < 0.0005, ANOVA). The data were expressed as mean ± SD; n = 3. **e** Representative TIRF microscopic images and their corresponding histograms of PD-L1 mRNA expression in single EVs derived from H1568 cells with/without IFN-γ stimulation in comparison to healthy donor EVs and PBS as controls. The single-EV PD-L1 mRNA signals were characterized with ^Au^SERP using PD-L1-targeting CLN-MBs. H1568 EVs were spiked in healthy donor EVs at a 1:1 ratio with 5 × 10^10^ particles/mL each. The images were cropped and enlarged from their original images, which are provided in Supplementary Fig. S4a. **f** A performance evaluation of ^Au^SERP for PD-L1 mRNA detection in comparison to qRT-PCR. EVs derived from IFN-γ-stimulated H1568 cells were spiked in healthy donor EVs at different concentrations ranging from 0 to 5 × 10^10^ particles/mL. The healthy donor EV concentration was kept constant at 5 × 10^10^ EVs/mL for all samples. The LOD of ^Au^SERP for PD-L1 mRNA detection was ~ 10^6^ spiked tumor EVs, ~ 1000 times lower than qRT-PCR. The data were expressed as mean ± SD; n = 3. TFI, total fluorescence intensity; a.u., arbitrary units.

To efficiently deliver MBs into EVs for RNA hybridization, we next encapsulated the designed MBs into CLNs. We previously reported that EV-encapsulated cargo RNAs could be detected using CLNs containing target-specific MBs (*39*). CLN-MBs revealed an “onion-like” structure with multiple wrapped lipid-MB-lipid layers (**Fig. 3c/i**), as previously described (*42, 50*). The negatively charged MBs act as a bridge between the two positively charged liposomes. This process was repeated to form many layers in a single particle. Due to their positive charge, CLNs could fuse with negatively charged EVs via electrostatic interactions to form larger complexes, as shown in our cryo-TEM image (**Fig. 3c/ii**). This fusion allowed efficient delivery of MBs into the EVs, leading to the binding of MBs to the target RNAs within the complexes’ nanoscale confinement. The specificity of PD-L1 CLN-MBs was examined using artificial EVs containing PD-L1 single-stranded DNA (ssDNA). The resulting fluorescence signal showed a strong linear relationship with the concentration of artificial EVs, spiked in healthy donor EVs ranging from 0 to 10% (R^2^ = 0.9665, *p* < 0.0005, **Fig. 3d**), indicating the high specificity of PD-L1 CLN-MBs to hybridize with PD-L1 mRNA cargo.

We further employed PD-L1 CLN-MBs for single-EV PD-L1 mRNA characterization with ^Au^SERP. Consistent with the single-EV PD-L1 protein characterization, ^Au^SERP successfully detected PD-L1 mRNA in the single EVs derived from H1568 cells and quantitatively differentiated between the single EVs originating from cells with and without IFN-γ stimulation (*p* < 0.005, **Fig. 3e & Supplementary Fig. S4a**). EVs produced from IFN-γ-stimulated H1568 cells were also spiked into healthy donor EVs to determine the LOD of our platform for single-EV mRNA characterization compared to conventional qRT-PCR. ^Au^SERP could detect as low as ~ 10^6^ spiked H1568 EVs, which exceeded the sensitivity afforded by qRT-PCR by ~ 1000 times (**Fig. 3f**). Moreover, our system enabled the multiplexed detection of PD-L1 protein and mRNA biomarkers simultaneously (**Supplementary Fig. S4b**).

### ^Au^SERP enables sensitive single-EV PD-1 protein and mRNA detection

T cells activated with anti-CD3/CD28 antibodies and Interleukin 2 (IL-2) have been known to stimulate PD-1 expression on the cells (*51*). EVs collected from activated T cells were purified and examined for PD-1 protein and mRNA expression with ^Au^SERP. The size distribution of the purified EVs is shown in **Supplementary Fig. S5a**. T cell EVs were spiked into healthy donor EVs at different concentrations ranging from 0 to 5 × 10^10^ particles/mL. The healthy donor EVs were kept constant at 5 × 10^10^ particles/mL. Single EVs with detectable PD-1 protein levels on their surface appeared as bright spots in TIRF images (**Supplementary Fig. S5b**). When the spiked EVs exceeded 10^6^, the PD-1 protein fluorescence intensities increased proportionally to the spiked EV concentration and were significantly higher than intensities from the healthy donor EVs and the blank control (*p* < 0.05, **Supplementary Fig. S5c**). Therefore, our assay allowed comparative and quantitative evaluations of PD-1 protein expression on single EVs. PD-1 mRNA in the single EVs was successfully detected and quantified with ^Au^SERP (**Supplementary Fig. S5d**). Taken together, we effectively developed a highly sensitive ^Au^SERP biochip to quantify four biomarkers (PD-1/PD-L1 proteins and mRNAs) on the surface and inside single EVs.

### ^Au^SERP enables single-EV PD-1/PD-L1 protein and mRNA detection from NSCLC patient serum

Fifty-four patients with NSCLC at stage IV were enrolled for this study regardless of their PD-L1 IHC results (**Supplementary Table S1**). Thirty-seven patients underwent anti-PD-1 therapy with nivolumab, sixteen patients underwent anti-PD-1 therapy with pembrolizumab, and one patient underwent anti-PD-L1 therapy with atezolizumab. We classified the patients who showed a partial response (PR) or stable disease (SD) lasting more than 6 months as “responders” (n = 27), and the patients who showed progressive disease (PD) as “non-responders” (n = 27). PR, SD, and PD were defined following the Response Evaluation Criteria in Solid Tumors (RECIST version 1.1) (*52*). The patients’ tissue and serum samples were collected before starting immunotherapy. Tissue samples were characterized using the Dako PD-L1 IHC 22C3 pharmDx platform. Tissue samples with tumor cells positive for PD-L1 protein expression equal to or greater than 1% were positive for PD-L1 IHC (**Fig. 4a**). Of the patients within the cohort, 29 demonstrated a positive PD-L1 IHC result, and 10 demonstrated a negative PD-L1 IHC result. Meanwhile, 15 patients were marked as unknown since their PD-L1 status was absent from their medical records, and their formalin-fixed paraffin-embedded samples were not available for testing (**Fig. 4b**).

**Fig. 4.**
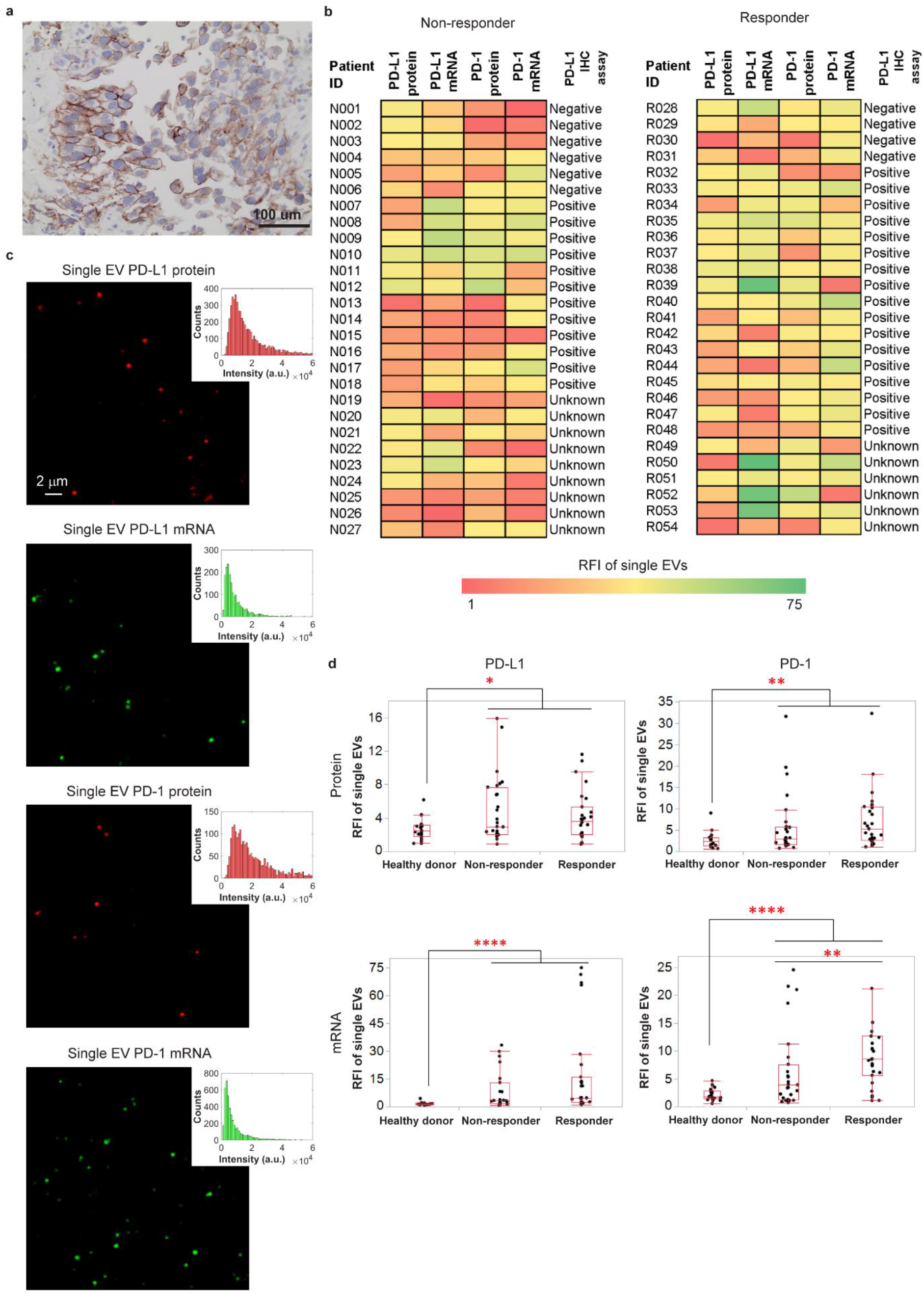
Measurements of PD-1/PD-L1 protein and mRNA biomarkers levels in single EVs from NSCLC patient serum with ^Au^SERP. **a** Representative IHC staining image of PD-L1 protein in tissue biopsies. PD-L1 protein was stained brown (by Dako PD-L1 IHC 22C3 pharmDx), and cell nuclei were stained blue (by hematoxylin). **b** Heatmap of single-EV PD-1/PD-L1 protein and mRNA expression levels measured with ^Au^SERP compared to the corresponding IHC score for PD-L1 expression in patient tissue (negative, positive, or unknown). **c** Representative TIRF microscopic images and their corresponding histograms of single-EV PD-1/PD-L1 protein and mRNA biomarkers characterized with ^Au^SERP. The images were cropped and enlarged from their original images, which are provided in Supplementary Fig. S6. **d** Box plots of quantitative fluorescence intensities of PD-1/PD-L1 protein and mRNA expression levels (\**p* < 0.05, \*\**p* < 0.01, \*\*\*\**p* < 0.0001, Mann-Whitney *U* test). RFI, relative fluorescence intensity; a.u., arbitrary units. 54 patients were evaluated (27 responders and 27 non-responders), along with 20 healthy donors.

EVs were purified from patient serum samples for characterization of single-EV PD-1/PD-L1 protein and mRNA biomarkers with ^Au^SERP. Using 3 μL of purified serum, all four biomarkers in the single EVs of the NSCLC patients were successfully detected with PD-1/PD-L1 antibodies and CLN-MBs. Representative TIRF images and histograms of the PD-1/PD-L1 protein and mRNA fluorescence signals at the single-EV level are shown in **Fig. 4c & Supplementary Fig. S6**. Single-EV PD-1/PD-L1 protein and mRNA expression levels of the NSCLC patients, in comparison with those of healthy donors, were quantified using the custom-built MATLAB algorithm (**Fig. 4d**). The capability of ^Au^SERP to detect single-EV protein and mRNA signals at such a small volume of serum suggests its potential applications as a non-invasive and sensitive approach to diagnose cancer patients and predict immunotherapy responses.

### ^Au^SERP enables single-EV characterization in subpopulations

^Au^SERP offers excellent flexibility in capturing and sorting EVs into subpopulations based on their membrane protein compositions by using the corresponding capture antibodies. Therefore, we wanted to know which EV subpopulation could provide the strongest PD-1/PD-L1 protein and mRNA signals to predict immunotherapy responses better. Given that PD-L1 can be highly expressed in tumor cells, a subpopulation of tumor-associated EVs was captured using anti-EGFR/EpCAM antibodies, which were shown to be efficient in capturing lung circulating tumor cells (CTCs) (*53, 54*). Anti-CD63/CD9 antibodies were also used to sort out the CD63^+^/CD9^+^ EV subpopulation. These two EV subpopulations derived from a small cohort of healthy donors, non-responders, and responders were examined for single-EV PD-L1 protein and mRNA signals. The CD63^+^/CD9^+^ EV subpopulation provided stronger PD-L1 signals with greater differentiation between samples (*p* < 0.05, **Supplementary Fig. S7a**). We also found high levels of PD-L1 protein expression in the cytoplasm of H1568 cells colocalizing with CD63 proteins (**Supplementary Fig. S7b**). In addition to tumor cells, PD-L1 is known to be robustly upregulated on tumor-infiltrating immune cells (macrophages, dendritic cells, and T cells) (*55, 56*). PD-L1 expression on these immune cells has pleiotropic effects on innate and adaptive immune tolerance in cancer. These may explain the greater efficacy of PD-L1 protein and mRNA detection in single EVs captured by anti-CD63/CD9 antibodies, since this cocktail captures EVs from all cellular sources, while anti-EGFR/EpCAM antibodies only capture EVs from tumor cells.

Given that PD-1 is highly expressed in activated T cells, T cell EVs subpopulations were captured using either anti-CD3, anti-CD4, anti-CD8, or anti-CD4/CD8 antibodies. With a small cohort of healthy donors, non-responders, and responders, we compared the single-EV PD-1 protein and mRNA signals. The CD63^+^/CD9^+^ EV subpopulation provided more consistent and sensitive PD-1 signals than the T cell-specific subpopulation (**Supplementary Fig. S8**). In addition to activated T cells, PD-1 can be expressed on natural killer cells, T cells, B cells, dendritic cells, and activated monocytes (*57*). Our results suggest that anti-CD63/CD9 antibodies that capture CD63^+^/CD9^+^ EVs from all cellular sources are preferable for single-EV PD-1/PD-L1 protein and mRNA characterization. Therefore, we used anti-CD63/CD9 antibodies to capture and sort EVs to detect all four biomarkers in single EVs for all NSCLC patient samples.

### ^Au^SERP offers an excellent single-EV characterization assay to diagnose NSCLC

We asked whether any single-EV single biomarkers (PD-1/PD-L1 proteins and mRNAs) could be good predictors for NSCLC diagnosis with ^Au^SERP. To address this question, EVs purified from serum samples of the 54 NSCLC patients (27 non-responders and 27 responders) and 20 healthy donors were characterized for all four biomarkers with ^Au^SERP. PD-1/PD-L1 protein and mRNA levels in patient samples (including responders and non-responders) were significantly higher than those of healthy donors (*p* < 0.05 for PD-L1 protein, *p* < 0.005 for PD-1 protein, *p* < 0.0001 for PD-1 mRNA and *p* < 0.00001 for PD-L1 mRNA, **Fig. 4d**). This suggests that our single-EV PD-1/PD-L1 protein and mRNA assays have potential in cancer diagnosis of NSCLC. Recent efforts in characterizing circulating EV PD-L1 protein have also shown that PD-L1 protein is a good biomarker to distinguish between cancer patients and healthy donors and can even determine the stage in cancer progression (*20, 58, 59*). A receiver operating characteristic (ROC) curve analysis (**Fig. 5a & Supplementary Fig. S9a**) demonstrated that compared to single-EV PD-L1 protein (area under the curve (AUC) = 0.669), the single-EV PD-L1 mRNA biomarker exhibited considerably higher accuracy in differentiation between NSCLC patients and healthy donors (AUC = 0.838), suggesting the single-EV PD-L1 mRNA as a better biomarker for NSCLC diagnosis. An AUC of 0.7 - 0.8 is considered acceptable in diagnostic test assessment, while 0.8 - 0.9 is excellent, and more than 0.9 is outstanding (*60*). The single-EV PD-L1 mRNA is, therefore, an excellent biomarker for NSCLC diagnosis.

**Fig. 5.**
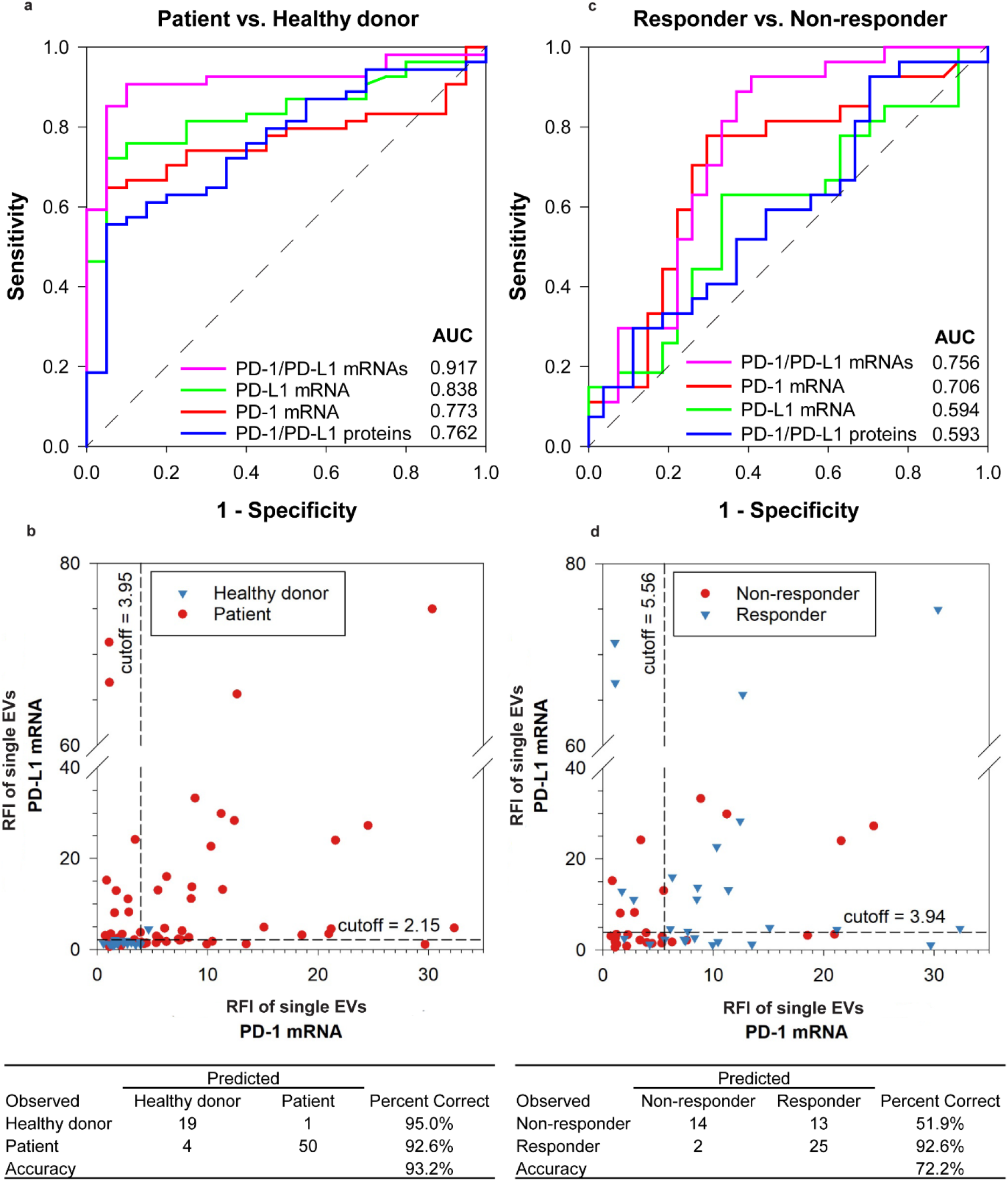
NSCLC diagnosis and prediction of immunotherapy response with ^Au^SERP. **a** Receiver operating characteristic (ROC) curves for NSCLC diagnosis based on single-EV analysis. **b** Scatter plot of fluorescence intensities of single-EV dual PD-1/PD-L1 mRNA biomarkers. Optimal cutoff values for NSCLC diagnosis based on single-EV PD-1/PD-L1 mRNA signals were obtained from the ROC curves in Fig. 5a. An individual is diagnosed with NSCLC if either the single-EV PD-1 mRNA signal is larger than 3.95 or if the single-EV PD-L1 mRNA signal is larger than 2.15, lending a diagnostic accuracy of 93.2%. **c** ROC curves for predictions of NSCLC patient responses to anti-PD-1/PD-L1 immunotherapy based on single-EV analysis. **d** Scatter plot of fluorescence intensities of single-EV dual PD-1/PD-L1 mRNA biomarkers. Optimal cutoff values for immunotherapy prediction based on single-EV PD-1/PD-L1 mRNA signals were obtained from the ROC curves in Fig. 5c. A patient is predicted as a responder if either the single-EV PD-1 mRNA signal is larger than 5.56 or if the single-EV PD-L1 mRNA signal is larger than 3.94, lending a prediction accuracy of 72.2%. 54 patients were evaluated (27 responders and 27 non-responders), along with 20 healthy donors.

Given the robustness of ^Au^SERP and its potential to co-quantify various single-EV protein and mRNA biomarkers, we hypothesized that a combination of the single-EV biomarkers (PD-1/PD-L1 proteins and mRNAs) would provide higher accuracy for NSCLC diagnosis. To test this hypothesis, a ROC curve analysis of different biomarker combinations was performed. Combining dual single-EV PD-1/PD-L1 mRNA biomarkers further increased the AUC to 0.917 (**Fig. 5a**), an outstanding level for cancer diagnosis. The data collected through ROC curve analyses were employed to determine cutoffs for the single-EV PD-1 mRNA signal at 3.95 or the single-EV PD-L1 mRNA signal at 2.15 (**Fig. 5b**). The accuracy of ^Au^SERP for this diagnosis was as high as 93.2%. This suggests the advantage of single-EV multi-biomarker analysis with ^Au^SERP. Therefore, we applied this multiplexed approach to predict patient responses to immunotherapy.

### ^Au^SERP single-EV dual PD-1/PD-L1 mRNA biomarkers analysis outperforms PD-L1 IHC at predicting responses to anti-PD-1/PD-L1 immunotherapy

To evaluate the performance of ^Au^SERP in predicting a patient’s response to immunotherapy, we compared single-EV PD-1/PD-L1 protein and mRNA signals from 54 NSCLC patients with their PD-L1 IHC assay results (**Fig. 4b**). Some responders demonstrated a positive PD-L1 IHC result and high single-EV PD-1/PDL1 protein and mRNA signals (e.g., patient R033, patient R035). Interestingly, for a responder with a negative PD-L1 IHC result (e.g., patient R028), the patient’s single-EV characterization with our platform indicated high PD-1/PD-L1 protein and mRNA signals. Furthermore, for a non-responder with a positive PD-L1 IHC score (e.g., patient N015), ^Au^SERP showed extremely low single-EV PD-1/PD-L1 protein and mRNA signals. These results suggest the potential of single-EV PD-1/PD-L1 proteins and mRNAs as better predictive biomarkers in comparison to PD-L1 IHC to anticipate patient responses to anti-PD-1/PD-L1 treatment.

For single-EV single biomarker analysis, there was no significant difference in PD-L1 protein and mRNA and PD-1 protein signals between responders and non-responders (*p* > 0.05, **Fig. 4d**). There were also no significant differences between the size distributions nor concentrations of EVs in serum samples of responders and non-responders (*p* > 0.05, **Supplementary Fig. S9b-c**). Single-EV PD-1 mRNA was the only biomarker with significantly higher levels in responders than in non-responders (*p* < 0.01, **Fig. 4d**). A ROC curve analysis also demonstrated that single-EV PD-1 mRNA had the highest AUC value (AUC = 0.706) among other single-EV single biomarkers (**Fig. 5c & Supplementary Fig. S9a**).

Interestingly, we found that mRNA biomarkers performed better for both PD-1 and PD-L1 than protein biomarkers as indicated by their higher AUC values (**Supplementary Fig. S9a**). The combination of single-EV dual PD-1/PD-L1 mRNA biomarkers further increased the AUC to 0.756, which was significantly higher than that of single-EV dual PD-1/PD-L1 protein biomarkers (AUC = 0.593, *p* < 0.05, **Fig. 5c**). Following the previously introduced cutoff approach, a patient could be predicted as a responder if either the single-EV PD-1 mRNA signal was larger than 5.56 or if the single-EV PD-L1 mRNA signal was larger than 3.94 (**Fig. 5d**). The accuracy of ^Au^SERP at predicting patient responses to immunotherapy was 72.2%, which is substantially higher than the commonly reported accuracy of the PD-L1 IHC assay at 20 ~ 40% (*61*). As such, single-EV dual PD-1/PD-L1 mRNA biomarkers characterized with ^Au^SERP outperformed the FDA-approved PD-L1 IHC assay at predicting responses to anti-PD-1/PD-L1 immunotherapy.

## DISCUSSION

NSCLC represents 80 – 85% of lung cancer, the second most common cancer and the leading cause of cancer-related death in both men and women in the U.S. (Cancer Statistics Center, American Cancer Society, 2020). Until now, the FDA has approved three ICIs targeting PD-L1 (atezolizumab, durvalumab, and avelumab) and two ICIs targeting PD-1 (nivolumab and pembrolizumab) for NSCLC patients. However, the objective response rates (ORR) to PD-L1 or PD-1 blockade remains 20 ~ 40%, even for patients with positive PD-L1 expression (*61*). This low ORR could be explained by the biology of PD-L1 expression within tumors, which is known to be both spatially and temporally variable. PD-L1 staining by IHC in one section at a given time does not reflect this variability and is referred to as a sampling error. Additionally, it is challenging to standardize PD-L1 IHC testing due to multiple scoring criteria and the inevitable subjective interpretation with pathologist scoring (*62*). Furthermore, using a single PD-L1 protein biomarker might not be sufficient to predict a reliable response to immunotherapy (*61*). EVs, shed from primary and metastatic tumors and circulate in the bloodstream (*63*), can better represent the tumor burden. Present in a non-invasive source, EVs can be readily collected at many time points before and after treatment to predict and surveil responses to immunotherapy. With our approach, single-EV protein and mRNA signals, shown as bright spots in TIRF images, can be accurately quantified using a custom-built algorithm that calculates the net fluorescence of every single EV, allowing for high-throughput analyses of biomarker levels. Therefore, issues of standardization and subjectivity can be avoided with ^Au^SERP. Moreover, ^Au^SERP offers a unique approach to quantify mRNA and protein biomarkers for a better cancer prognosis on immunotherapy.

PD-L1 and PD-1 are cellular transmembrane proteins secreted as EV proteins or freely soluble proteins (*64, 65*). In patients with cancer, extracellular PD-L1 protein may serve as an agent of immune suppression by inhibiting T cell activation and/or counteracting biomolecules for immune activation (*20, 47, 64*). Meanwhile, extracellular PD-1 protein plays an adjuvant role in enhancing antigen-specific T cell immunity responses (*65, 66*). Previous studies have shown that PD-L1 proteins are present EVs surfaces isolated from plasma/sera of patients with metastatic melanoma (*20*), head and neck squamous cell carcinoma (*21*), glioma (*67*), and NSCLC (*58, 68*). In addition to PD-L1 proteins, PD-L1 mRNAs have been demonstrated to exist in EVs derived from saliva and plasma of patients with periodontitis and melanoma/NSCLC, respectively (*22, 23*). Compared to EV PD-L1 protein/mRNA, studies on EV PD-1 protein/mRNA are limited (*21*). Furthermore, most of these studies used western blot, magnetic bead-based flow cytometry, RT-PCR, or ddPCR to measure EV protein and mRNA, which are bulk-analysis methods with limited sensitivity in capturing the heterogeneity present in EVs. The technology we developed offers a highly sensitive method to quantify molecular contents in EVs at the single-EV level by combining ^Au^SERP with TIRF microscopy to simultaneously measure PD-1/PD-L1 protein and mRNA contents on the same device.

The presence of 30-nm gold NPs significantly enhanced the sensitivity of single-EV analyses with ^Au^SERP. Metal nanoparticles exhibit localized SPR, which can be seen by a strong UV-Vis absorption band that is absent from bulk metal (*69*). Gold NPs are better than other NPs in signal enhancement since gold NPs can cause higher RI changes and resonance angle shifts (*46*). Gold spherical NPs have been widely used to amplify the signal and improve the sensitivity of SPR biosensors (*70–72*). The LOD of prostate-specific antigens from 10 ng/mL to sub ng/ml range was improved by introducing streptavidin-conjugated gold NPs to a biotinylated secondary antibody (*73*). In cancer biomarker detection using SPR biosensors, 20-nm gold NPs reduced the LOD 8 times, while 40-nm gold NPs lowered the LOD 65 times in comparison to the sandwich assay (*46*). By measuring the SPR angle shift in the presence of aqueous methanol, 30 nm was found to be the optimal particle size when compared to 10 and 60 nm NPs (*74*). In another study, four different sizes were tested (13, 30, 40, and 50 nm) to develop a highly sensitive nuclease-linked fluorescence oligonucleotide assay in which the 40-nm gold NPs yielded the strongest signal (*75*). As can be observed, the optimal NP size varies by device. For ^Au^SERP, the optimal NP size was 30 nm.

By combining PEG, BSA, and Tween-20 to block non-specific bindings, TIRF microscopy for high-resolution imaging, a thin gold coating and gold NPs for SPR signal enhancement, and amplification methods for detection by antibodies (using TSA) and MBs (using CLNs), we engineered a highly sensitive platform. ^Au^SERP exhibited a significantly high SNR of 482.02 ± 21.81, which enabled us to accurately quantify low expression biomarkers such as PD-1/PD-L1 proteins and mRNAs at a single-EV level. Our platform was ~ 1000 times more sensitive than ELISA and qRT-PCR in quantifying PD-L1 protein and mRNA, respectively. With such high sensitivity, our technology only required ~ 3 μL of purified serum for protein and mRNA biomarker characterization. ^Au^SERP is also capable of high-throughput analyses with up to 256 individual samples per assay. In addition to being a sensitive and high-throughput assay, ^Au^SERP enabled multiplex detection of different protein and mRNA biomarkers simultaneously. Moreover, our platform offered great flexibility in capturing and sorting EVs into subpopulations to study the function of each EV subtype.

EV PD-L1 protein was previously reported as a potential predictive biomarker to distinguish patients with metastatic melanoma who respond to anti-PD-1 therapy (*20*). Elevated levels of pre-treatment EV PD-L1 following a ceased increase in on-treatment PD-L1 levels were observed in non-responders, which may reflect the exhaustion of T cells. Meanwhile, responders displayed a larger increase in EV PD-L1 levels as early as 3–6 weeks following the initial treatment, which correlates positively with T cell reinvigoration (*20*). However, for NSCLC, identifying responders from non-responders using liquid biopsies is still very challenging (*64*). With ^Au^SERP, we successfully demonstrated a comprehensive profile of four immunotherapy biomarkers (PD-1/PD-L1 proteins and mRNAs) at a single-EV resolution from EVs isolated from the serum of NSCLC patients. A cohort of 27 non-responders and 27 responders was examined in our study. Interestingly, a ROC curve analysis of four single-EV biomarkers suggests that single-EV mRNAs are better than single-EV proteins for both cancer diagnosis and immunotherapy response predictions. This could be due to the ability of EVs to preserve and protect mRNAs within their phospholipid membrane from degradation *in vivo*, while EV surface proteins are exposed to protease degradation. ^Au^SERP offers a robust and highly sensitive approach to characterize single-EV mRNA biomarkers in which single EVs are captured and directly detected using CLN-MBs without the need for tedious RNA extraction, cDNA reverse transcription, and qRT-PCR. The dual single-EV PD-1/PD-L1 mRNA biomarkers achieved AUC values of 0.917 and 0.756 to distinguish patients from healthy donors and responders from non-responders. The accuracy of ^Au^SERP to predict patient responses to immunotherapy was 72.2%, which exceeded the FDA-approved PD-L1 IHC assay. Our study, therefore, showed that pre-treatment single-EV dual PD-1/PD-L1 mRNA are good predictors to identify NSCLC patients that will benefit from anti-PD-1/PD-L1 immunotherapy. Given the heterogeneity and dynamic changes of PD-1/PD-L1 expression in tumors and the invasive nature of tumor biopsies, developing a non-invasive single-EV assay with ^Au^SERP is an attractive alternative to IHC scoring. Longitudinal assays before and shortly after administration of ICIs would be an exciting avenue to explore to improve prediction accuracy.

A patient-oriented approach to immunotherapy using predictive biomarkers is desired to maximize clinical benefit, improve cost-effectiveness, and reduce the economic burden of the disease. A previous study showed that compared to treating all NSCLC patients with anti-PD-1/PD-L1 immunotherapy, the selection of patients based on positive PD-L1 IHC scores improved incremental quality-adjusted life years by up to 183% and decreased the incremental cost-effectiveness ratio by up to 65% (*76*). However, due to its limited precision, there are scenarios where patients with tumors scored as PD-L1-positive do not respond to anti-PD-1/PD-L1 therapy (false-positive testing) and *vice versa (61*). In the case of false-negative testing, it could diminish the number of potential life-years saved. The success of this work in offering a more accurate prediction of NSCLC patient responses can, therefore, improve the survival of patients and minimize the cost of treatment. Furthermore, the success ^Au^SERP is a breakthrough in cancer therapy in which personalized cancer immunotherapy can be achieved by feasibly identifying patients most likely to benefit from immunotherapy and monitoring their responses throughout their treatment. In principle, our ^Au^SERP technology applies to a broad spectrum of biomedical applications (e.g., early cancer diagnosis, neurodegenerative diseases such as Alzheimer’s, Traumatic Brain Injury, viral diseases, and cardiovascular diseases), in which any combination of antibodies and MBs could be used to detect disease-specific proteins and RNAs of interest in specific membrane-enveloped subpopulations.

## MATERIALS AND METHODS

### ^Au^SERP fabrication

A high precision glass coverslip (#D 263 M Glass, 24 × 75 mm rectangle, 0.15 mm thickness Schott AG, Mainz, Germany) was cleaned with deionized (DI) water and ethanol two times alternately in an ultrasonic bath for 5 min each and then dried with nitrogen gas. Subsequently, the cleaned glass coverslip was activated using a UV-ozone cleaner (UVO Cleaner Model 42, Jelight, Irvine, CA) for 15 min. Thin layers of 2-nm-thick Ti and 10-nm-thick Au were then deposited, respectively, using a Denton e-beam evaporator (DV-502A, Moorsetown, NJ). After Au deposition, the freshly prepared Au-coated glass was immersed into a linker solution containing β-mercaptoethanol (βME, Sigma-Aldrich, St. Louis, MO), PEG-SH (#MPEG-SH-2000, Laysan Bio Arab, AL), and biotin-PEG-SH (#PG2-BNTH-2k, Nanocs, New York, NY) (molar ratio = 95:3:2) in 200 proof ethanol (Fisher Scientific, Hampton, NH) for overnight in the dark at 22°C. The glass coverslip was then rinsed with ethanol to remove any excess mixtures and subsequently air-dried. The treated glass was attached to a 64-well gasket (Grace Bio-Labs ProPlate tray set, Sigma-Aldrich) and washed thoroughly with DI water. Next, 0.005% (w/v) streptavidin-conjugated gold nanoparticles (NPs, Nanocs) in PBS were introduced into the wells for 2 h at room temperature (RT) on a rocker (24 rpm, Benchmark Scientific, Sayreville, NJ). Different sized NPs (5, 30, and 50 nm) were used to test the EV capture efficacy and non-specific binding of antibodies. After rinsing three times with PBS, the surface was incubated with a capture antibody cocktail for 1 hr at RT on the rocker. The cocktail included 10 μg/mL each of a mouse anti-CD63 monoclonal antibody (#MAB5048, R&D Systems, Minneapolis, MN) and a mouse anti-CD9 monoclonal antibody (#MAB1880, R&D Systems). These antibodies were biotinylated using an EZ-Link micro Sulfo-NHS-biotinylation kit (ThermoFisher Scientific, Waltham, MA) before the incubation. After the antibodies were tethered onto the gold surface, the free antibodies were washed away three times with PBS, and then blocked with 3% (w/v) BSA (Sigma-Aldrich) and 0.05% (v/v) Tween 20 (Sigma-Aldrich) in PBS for 1 h at RT before EV capture.

### Atomic force microscopy (AFM)

The topography of the surfaces coated with different sized streptavidin-conjugated gold NPs was characterized using an AFM (Asylum Research MFP-3D-BIO AFM, Oxford Instruments, Abingdon, United Kingdom). Before imaging, the surfaces were rinsed thoroughly with DI water to avoid salt crystals and air-dried.

### Cell Culture

H1568 cells (NCI-H1568, ATCC CRL-5876, Manassas, VA) were cultured in a growth medium containing RPMI 1640 (ThermoFisher Scientific), 10% (v/v) fetal bovine serum (FBS, Sigma-Aldrich), and 1% (v/v) penicillin-streptomycin (PS, ThermoFisher Scientific). The medium was replaced every 2 to 3 days, and cultures were maintained in a humidified incubator at 37°C with 5% CO_2_. When the cells reached 80%–90% confluence, they were detached using TrypLE express enzyme (ThermoFisher Scientific) and passaged at 1:3–1:6 ratios. H1568 cells at passages of 6-10 were used in this study.

To stimulate PD-L1 expression, H1568 cells were first grown to 80% confluency in the growth medium in a cell culture flask (Corning Inc., Corning, NY). The cells were then washed with PBS and changed to an RPMI medium supplemented with 10% (v/v) EV-depleted FBS, 1% (v/v) PS, and 100 ng/ml recombinant human IFN-γ (Peprotech, Rocky Hill, NJ) for 48 hr. The cells cultured in the medium without IFN-γ supplements were used as controls. EV-depleted FBS was collected from the filtrate from FBS introduced through tangential flow filtration (TFF) system with a 500 kDa molecular weight cutoff (MWCO) hollow fiber filter (polysulfone, Repligen, Waltham, MA). After 48 h, the culture supernatant was centrifuged at 2000 × g for 10 min (Centrifuge 5810R, Eppendorf, Hauppauge, NY) to remove cell debris before EV purification. On the other hand, the adherent cells in the flasks were detached and then pelleted by centrifugation at 1000 rpm for 5 min. The cell pellets were used to characterize the PD-L1 protein and mRNA in the cells using IHC staining, ELISA, and qRT-PCR as described in the following sections.

### EV purification from cell culture supernatants and clinical samples

Healthy donor and NSCLC patient blood samples were obtained with appropriate informed consent under approved Institutional Review Board (IRB) protocols #2018H0268 and #2015C0157, respectively, at The Ohio State University. The study was carried out following relevant guidelines and regulations. Blood samples from stage IV NSCLC patients were collected before they underwent immunotherapy. Serum was separated from blood using a BD Vacutainer Serum Separation Tube (SST, #367985, Becton Dickinson) according to the manufacturer’s protocol.

The prepared cell supernatants and sera were first filtered through 1-μm filters (GE Healthcare Whatman Puradisc GMF, Fisher Scientific). They were subsequently concentrated and diafiltrated using TFF with a 500 kDa filter for purification as described (*77*). After TFF, the retentates were concentrated using centrifugal units (10 kDa MWCO, MilliporeSigma Amicon Ultra Centrifugal Filter Unit, Fisher Scientific) at 3000 × g for 20 min. The concentration of EVs was quantified using a tunable resistive pulse sensing (TRPS) technology (qNano Gold instrument, Izon Science, Medford, MA) with NP150 (target size range 70 – 420 nm) and NP600 (target size range 275 – 1570 nm) nanopore membranes.

### CD63 detection of EVs

EVs harvested from H1568 cells without IFN-γ stimulation were adjusted to a concentration of 10^10^ particles/mL. After that, the purified EVs were added onto ^Au^SERP biochips coated with different NP sizes. PBS was used as a blank control. The following incubation and washing steps were performed at RT on a rocker. EVs were captured for 2 h, washed three times with PBS, and then blocked with 3% (w/v) BSA and 0.05% (v/v) Tween 20 in PBS for 1 h. The samples were subsequently incubated with a mouse anti-CD63 monoclonal antibody (MX-49.129.5) – Alexa Fluor 488 conjugate (#sc-5275 AF488, Santa Cruz Biotechnology) at a dilution of 1:200 in 1% (w/v) BSA in PBS for 1 h. Next, the samples were rinsed three times with 0.05% (v/v) Tween 20 in PBS. Images were taken using a TIRF microscope (Nikon Eclipse Ti Inverted Microscope System). The images were recorded by an Andor iXon EMCCD camera with a 100x oil lens at the same laser power and exposure time. For each sample, 100 images (10 × 10 array) were collected.

### PD-L1 protein staining of cells and tissues

The cell pellets were fixed in a 4% formaldehyde solution in PBS (Fisher Scientific) for 2 h at RT and then washed with PBS by centrifugation. The pellets were subsequently blocked in agarose gel and embedded in paraffin. They were then sectioned into 5-μm thick slices, stained for PD-L1 protein, and then counterstained with hematoxylin using an automated Dako PD-L1 IHC 22C3 pharmDx platform (Agilent Technologies, Santa Clara, CA). The images were taken using a light microscope (CKX41 Inverted Microscope, Olympus, Tokyo, Japan).

Tissue biopsies were obtained with informed consent from NSCLC patients using the approved IRB protocol #2015C0157 at The Ohio State University. Tissue and blood samples were collected from each patient before starting immunotherapy. Classification of responders or non-responders to immunotherapy and PD-L1 IHC results were obtained from medical reports. PD-L1 IHC of the tissue biopsies were performed using the automated Dako PD-L1 IHC 22C3 pharmDx platform.

### PD-1/PD-L1 protein detection of EVs

The purified EVs from H1568 cells (with and without IFN-γ stimulation) and blood samples (healthy donors and NSCLC patients) were captured with ^Au^SERP. PBS was used as a blank control, and a washing buffer. All incubation and washing steps were conducted at RT on the rocker. After capture, the samples were rinsed three times and stained for PD-1/PD-L1 proteins using an Alexa Fluor 647 Tyramide SuperBoost kit (#B40926, ThermoFisher Scientific). Firstly, the EVs were fixed with 4% formaldehyde solution in PBS for 10 min. After washing, a 3% hydrogen peroxide solution was added to quench the endogenous peroxidase activity of the samples for 15 min, followed by incubation with 3% (w/v) BSA and 0.05% (v/v) Tween 20 in PBS for 1 h. Rabbit anti-PD-L1 monoclonal antibody (#86744S, Cell Signaling Technology, Danvers, MA) or rabbit anti-PD-1 monoclonal antibody (#86163S, Cell Signaling Technology) was then diluted 500-fold in a blocking buffer and incubated for 1 h. Next, the samples were washed three times for 10 min each before incubation with a poly-HRP-conjugated secondary antibody for 1 h. After washing three times for 10 min each, a tyramide working solution was applied for 10 min. The reaction was stopped using a stop reagent. After that, the samples were rinsed three times and imaged using the TIRF microscope as mentioned above.

### Design and fabrication of CLN-MBs

MBs (listed 5′-3′) targeting PD-1 and PD-L1 mRNAs used in this study were +GGT +CCT /iCy3/ +CCT +TCA +GGG GCT GGC GCC CCT GAA GG /BHQ_2/ and +GGT +AGC /iCy3/ +CCT +CAG +CCT GAC ATG AGG CTG AGG /BHQ_2/, respectively. They were designed based on an NCBI reference sequence of PD-1 (NM_005018.3) and PD-L1 (NM_014143.4) using Primer3 and BLAST (Primer-BLAST) provided by NCBI-NIH. Locked nucleic acid nucleotides (positive sign (+) bases) were incorporated into oligonucleotide strands to improve the thermal stability and nuclease resistance of MBs for incubation at 37°C. The designed MBs were custom synthesized and purified by Integrated DNA Technologies (IDT, Coralville, IA).

To fabricate CLN-MBs, an aqueous solution of MBs in PBS was vigorously mixed with a lipid formulation of 1,2-dioleoyl-3-trimethylammonium-propane (DOTAP, Avanti Polar Lipids, AL, USA), Cholesterol (Sigma-Aldrich), 1,2-dioleoyl-sn-glycero-3-phosphocholine (DOPC, Avanti Polar Lipids), and Bis(1,2-distearoyl-sn-glycero-3-phosphoethanolamine)-N-[(polyethylene glycol)-2000] (Bis-DSPE PEG2000, Avanti Polar Lipids) (50:30:18:2 mole ratio in 200 proof ethanol), and then sonicated for 5 min using an ultrasonic bath. The MB/lipid mixture was subsequently injected into PBS, vortexed, and sonicated for 5 min. Finally, the mixture was dialyzed with a 20 kDa MWCO to remove free MBs.

### mRNA staining of cells

H1568 cells were seeded at a density of 10^5^ cells/mL in 16-well chambers (Grace Bio-Labs ProPlate tray set) attached to a glass slide (Fisher Scientific). To stimulate PD-L1 expression, the cells were incubated with 100 ng/ml recombinant human IFN-γ in the growth medium for 48 h. Cells without IFN-γ stimulation were employed as a control. The H1568 cells were then fixed in 4% formaldehyde solution in PBS for 30 min at RT and then permeabilized with 0.2% (v/v) Triton X-100 (Sigma-Aldrich) in PBS for 10 min at RT. The cells were subsequently rinsed with PBS and incubated with 0.5 μM free PD-L1 MBs for 4 h at 37°C. After washing three times with PBS for 5 min each, the glass slide was detached and mounted onto a cover glass (Fisher Scientific) using ProLong Gold Antifade Mountant with DAPI (ThermoFisher Scientific). The images were taken using a confocal microscope (Olympus FV3000).

### Calibration of CLN-MBs using artificial EVs

PD-L1 ssDNA oligo (listed 5′–3′) used in this study was CTG ACA TGT CAG GCT GAG GGC TAC CCC AAG. The designed oligo was custom synthesized and purified by IDT. To fabricate artificial EVs, an aqueous solution of the PD-L1 ssDNA oligo was mixed with a lipid formulation of 1,2-dioleoyl-sn-glycero-3-phosphoethanolamine (DOPE, Avanti Polar Lipids), linoleic acid (LA, Sigma-Aldrich), 1,2-dimyristoyl-rac-glycero-3-methoxypolyethylene glycol-2000 (DMG-PEG 2000, Avanti Polar Lipids) (50:48:2 mole ratio in 200 proof ethanol). The mixture was subsequently sonicated for 5 min, injected into PBS, and then sonicated for another 5 min. After dialysis with a 20 kDa MWCO to remove free molecules, the suspension was spiked into healthy donor EVs at different concentrations of 0.625, 1.25, 2.5, 5, and 10%.

For the calibration experiment, the gold-coated biochip was tethered with biotinylated PD-L1 CLN-MBs, which was made by replacing Bis-DSPE PEG2000 with DSPE-PEG(2000) Biotin (Avanti Polar Lipids) in the lipid formulation of CLN-MB fabrication. The artificial EVs at different concentrations were then captured on the biochip by fusion with the PD-L1 CLN-MBs for 2 h at 37°C. After rinsing with PBS, the samples were imaged using the TIRF microscope.

### PD-1/PD-L1 mRNA detection of EVs

The purified EVs from serum samples of healthy donors and cancer patients were captured with ^Au^SERP for 2 h at RT. After washing with PBS, PD-1/PD-L1 CLN-MBs were applied and incubated for 2 h at 37°C. The samples were rinsed with PBS and imaged using the TIRF microscope.

### Cryogenic transmission electron microscopy (cryo-TEM)

Cryo-TEM was used to characterize purified EVs from IFN-γ-stimulated H1568 cells, their PD-L1 proteins, PD-L1 CLN-MBs, and complexes of CLN-MBs fused with EVs. A concentration of 10^13^ EVs/mL was necessary for this experiment. The presence of PD-L1 protein on the surface membranes of EVs was verified using immunogold labeling. Goat PD-L1 polyclonal antibody (#AF156, R&D Systems) was firstly conjugated with gold NPs using a gold conjugation kit (#ab201808, Abcam, Cambridge, United Kingdom) according to the manufacturer’s instructions, and then incubated with the EVs for 1 h at RT. PD-L1 CLN-MBs were incubated with the EVs for 1 h at 37°C to observe their fusion.

Cryo-TEM samples were prepared by applying a small aliquot (3 μL) of the samples to a specimen grid. After blotting away excess liquid, the grid was immediately plunged into liquid ethane to rapidly form a thin layer of amorphous ice using a Thermo Scientific Vitrobot Mark IV system. The grid was transferred under liquid nitrogen to a Thermo Scientific Glacios CryoTEM. Images were recorded by a Thermo Scientific Felcon direct electron detector.

### Image analysis

All the spots in the TIRF image were first located with distinguished edges and background noise was removed by a Wavelet denoising method using a custom-built Matlab algorithm. The net fluorescence intensity of the spot was then calculated by summing the intensities of all the pixels within the spot, which were subtracted by the mean intensity of pixels surrounding the spot. Subsequently, a histogram of net fluorescence intensities of all the spots was obtained and their TFI was also calculated. All spots falling outside a threshold set through a user interface were not considered in the calculation. The relative fluorescence intensity (RFI) of an EV sample was defined as the ratio of the TFIEV to the TFI_neg_ within the same ^Au^SERP biochip.

### ELISA

PD-L1 protein expression levels in H1568 cells (with and without IFN-γ stimulation) and on the surface of H1568 EVs (with IFN-γ stimulation) were quantified using a PD-L1 Human ELISA kit (#BMS2212, ThermoFisher Scientific). For the cells, their pellets were lysed in RIPA buffer (ThermoFisher Scientific) with the addition of Thermo Scientific Halt Protease and Phosphatase Inhibitor Cocktails on ice for 5 min then centrifuged at 14,000 × g for 15 min to remove cell debris. For the EVs, they were spiked in healthy donor serum at different concentrations ranging from 0 to 10^11^ particles/mL. All the samples were subsequently incubated in the ELISA plate, and their PD-L1 expressions were quantitatively detected according to the manufacturer’s instructions. The PD-L1 concentration in the cell lysis was normalized to its total protein concentration, which was measured using a Pierce Rapid Gold BCA Protein Assay kit (ThermoFisher Scientific).

### qRT-PCR

PD-L1 mRNA levels in H1568 cells (with and without IFN-γ stimulation) and H1568 EVs (with IFN-γ stimulation) were quantified using qRT-PCR. Total RNA from the cells and EVs were first isolated and purified using an RNeasy Mini Kit and a miRNeasy Serum/Plasma kit (Qiagen, Hilden, Germany), respectively, according to the manufacturer’s instructions. cDNA was then synthesized from the total RNA using a High-Capacity cDNA Reverse Transcription kit (Applied Biosystems, Foster City, CA) on a thermal cycler (Veriti 96-Well Thermal Cycler, Applied Biosystems). Subsequently, PD-L1 mRNA expression was quantified using a TaqMan Gene Expression assay (ThermoFisher Scientific, Assay Id: Hs01125301_m1) on a Real-Time PCR System (Applied Biosystems). The cellular PD-L1 mRNA expression was normalized to GAPDH (Assay Id: Hs02758991_g1), a housekeeping gene.

### Statistical analysis

All *in vitro* experiments and assays were repeated three times. All clinical samples were measured two times. JMP Pro 14, MATLAB R2019a, Sigma Plot 14.0, and IBM SPSS Statistics 25 were used for data analysis. For *in vitro* samples, a significance of mean differences was determined using a two-tailed paired Student’s *t*-test. For clinical samples, a non-parametric Mann–Whitney *U*-test was used for analysis. Error bars shown in graphical data represent mean ± SD. A *p-*value below 0.05 was considered a statistically significant difference. The performance of classification schemes was evaluated using ROC curve analyses. For single biomarker analyses, ROC curves were determined from the expression level of each biomarker. For multiple biomarker analyses, ROC curves were plotted following a binary logistic regression. Optimal cutoff points were obtained from the ROC curves using the ROC curve analysis module in Sigma Plot. The accuracy of an assay was defined as

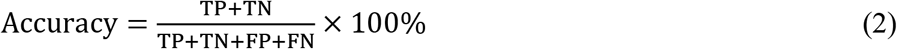

where TP, TN, FP, and FN represent the number of true positives, true negatives, false positives, and false negatives, respectively.

## Acknowledgments

We acknowledge all the patients and healthy volunteers who participated in this study. We thank Dr. Nick Chiang for his support with real-time PCR. We thank Dr. Xiaokui Mo for helpful advice on the statistical tests. Electron microscopy was performed at the Center for Electron Microscopy and Analysis (CEMAS) at The Ohio State University.

## Author contributions

L.T.H.N. prepared the figures and wrote the manuscript with input from all authors. L.T.H.N., K.J.K, L.J.L., and E.R. developed the technology. L.T.H.N., L.J.L., and E.R. designed the study and analyzed the data. L.T.H.N. and K.J.K. optimized and fabricated the ^Au^SERP biochip. L.T.H.N. performed cell culture experiments, ELISA, qRT-PCR, AFM, cryo-TEM, and biomarker characterization. K.J.K. designed the MBs. K.J.K. and X.W. fabricated the CLN-MBs. X.W., D.B. and X.Y.R. developed the custom-built algorithm for image analysis. L.T.H.N., J.Z., and M.J.Y. purified EVs using TFF. J.Z., M.J.Y, and X.Y.R. performed TRPS measurement. N.W. obtained healthy donor blood and isolated T cells. T.O., J.A., T.S., and D.C. provided clinical samples, PD-L1 IHC results, and patient clinical information. Y.M. performed confocal fluorescence imaging and assisted with qRT-PCR. X.Y.R., H.L., and A.P. assisted with ^Au^SERP preparation, EV purification, and manuscript writing. E.R. supervised the project. All authors have read and approved the final manuscript.

## Data and materials availability

All data needed to evaluate the conclusions in the paper are present in the paper and/or the Supplementary Materials. Additional data related to this paper may be requested from the authors.

