## Supplementary for "An Immunogold Single Extracellular Vesicular RNA and Protein (^Au^SERP) Biochip to Predict Responses to Immunotherapy in Non-Small Cell Lung Cancer Patients"

Table S1. Clinical characteristics of stage IV NSCLC patients.........................................16

Table S2. Antibodies used for single-EV capture and detection ................................................... 18

### SUPPORTING INFORMATION

#### Isolation and activation of T cells

Healthy donor blood was obtained with appropriate informed consent under an approved Institutional Review Board (IRB) protocol #2018H0268 at The Ohio State University. Briefly, blood was collected in a BD Vacutainer Plastic Blood Collection Tube with $K_2$EDTA (#366643, Becton Dickinson, Franklin Lakes, NJ) and lysed with red blood cell lysis buffer (150 mM ammonium chloride, 10 mM sodium bicarbonate, 0.1 mM EDTA, pH = 7.4) for 5 min at room temperature (RT). After centrifugation, a pellet was resuspended in phosphate buffer saline (PBS) and T cells were isolated using an immunomagnetic negative selection kit (EasySep Human CD8+ T Cell Enrichment Kit, #19053, Stemcell Technologies, Vancouver, Canada) according to the manufacturer's instructions. The cells were subsequently activated with ImmunoCult Human CD3/CD28 T Cell Activator (1:40 dilution, #10971, Stemcell Technologies) and Interleukin 2 (10ng/mL, Human Recombinant IL-2 (CHO-expressed), Stemcell Technologies) in a serum-free and xeno-free medium (ImmunoCult-XF T Cell Expansion Medium, Stemcell Technologies) for 3 days to stimulate PD-1 expression. After the incubation period, the culture was centrifuged at 2000 × g for 10 min to remove suspended cells and cell debris before EV purification.

#### Immunofluorescence staining of cells

H1568 cells grown on a glass slide in 16-well chambers were first fixed in a 4% formaldehyde solution in PBS for 15 min. Non-specific bindings were blocked with 3% (w/v) bovine serum albumin (BSA) and 0.05% (v/v) Tween 20 in PBS for 1 h at RT. A mouse anti-CD63 monoclonal antibody (MX-49.129.5) – Alexa Fluor 488 conjugate (#sc-5275 AF488, Santa Cruz Biotechnology), a mouse anti-CD9 monoclonal antibody (C-4) – Alexa Fluor 546 conjugate (#sc-13118 AF546, Santa Cruz Biotechnology), and a rabbit anti-PD-L1 monoclonal antibody – Alexa Fluor 647 conjugate (#41726S, Cell Signaling Technology) were diluted in 1% (w/v) BSA and 0.05% (v/v) Tween 20 in PBS and incubated with the cells for 1 hr at RT. After washing three times with PBS for 5 min each, the glass slide was detached and mounted onto a cover glass using ProLong Gold Antifade Mountant with DAPI. The images were taken with a confocal microscope (Olympus FV3000).

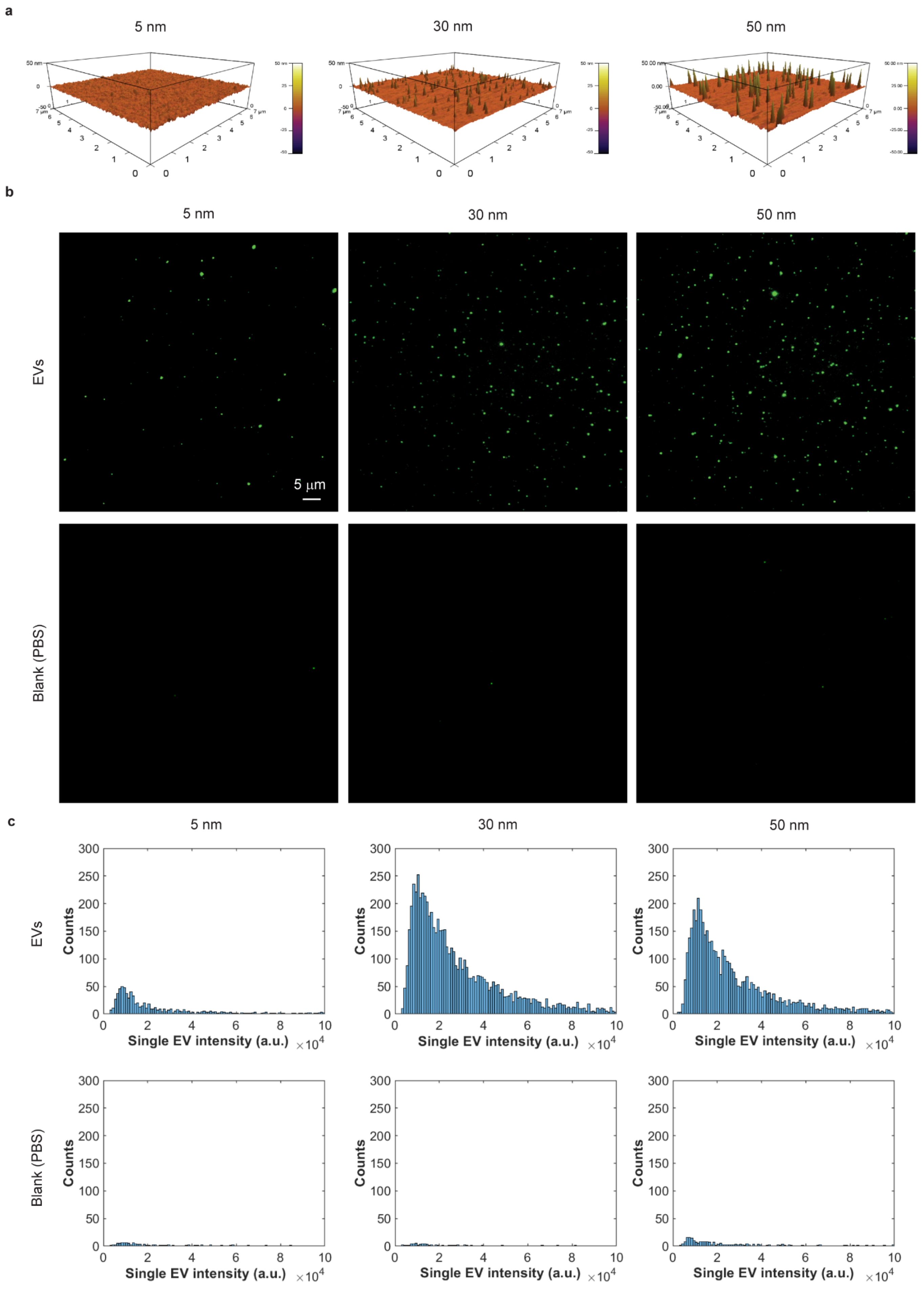

a
5 nm
30 nm
50 nm
b
5 nm
30 nm
50 nm
EVs
Blank (PBS)
5 μm
c
5 nm
30 nm
50 nm
EVs
Blank (PBS)
Counts
Single EV intensity (a.u.)
×10^4

**Fig. S1.** Characterization of $^{Au}$SERP coated with different-sized gold nanoparticles (5, 30, and 50 nm). **a** 3D height atomic force microscopy (AFM) images. **b** Representative total internal reflection fluorescence (TIRF) microscopic images of CD63 protein expression on the surface of H1568 single extracellular vesicles (EVs) in comparison with blank controls (PBS). **c** Distributions of fluorescence intensity of CD63 protein signals on single EVs. a.u., arbitrary units.

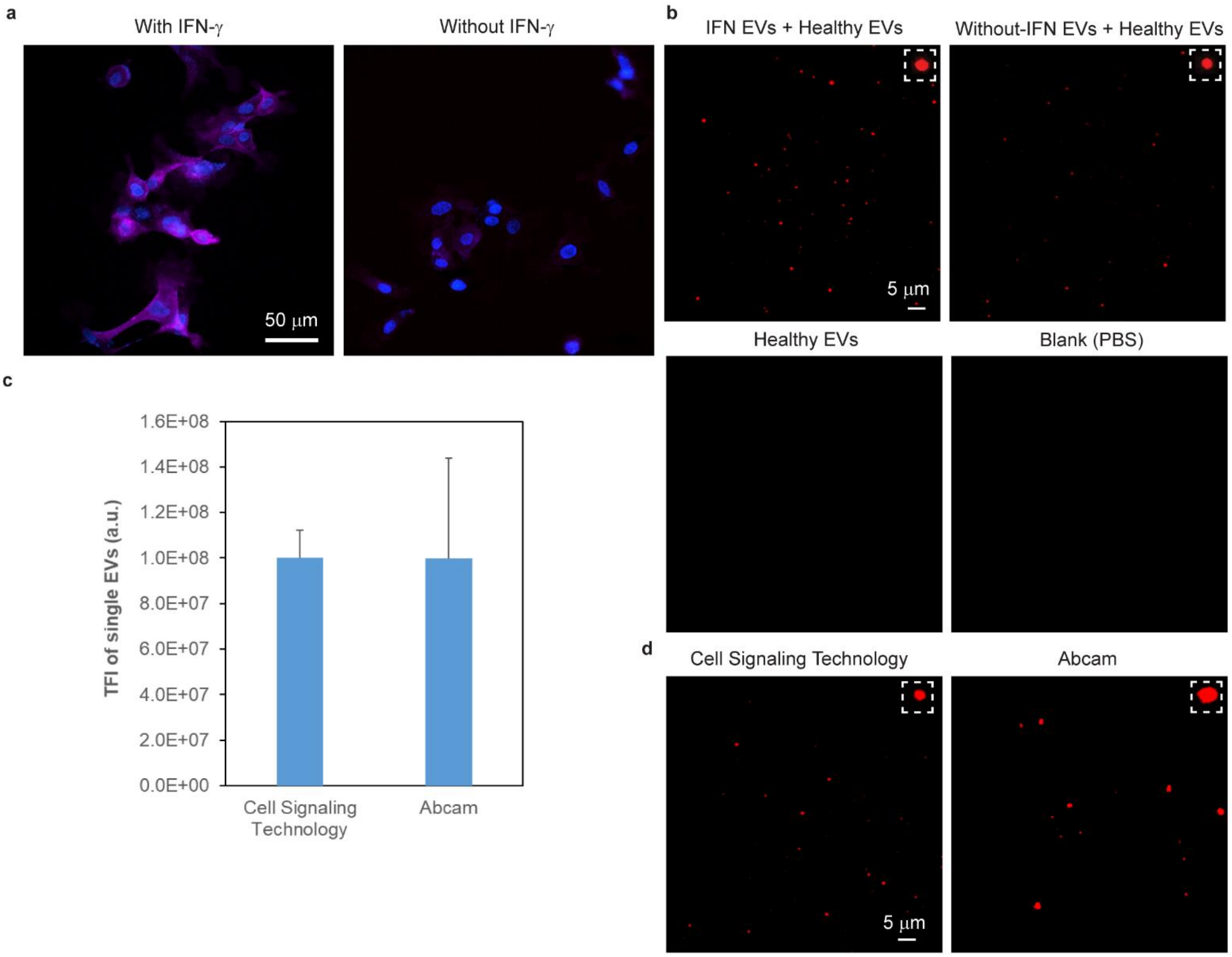

**Fig. S2.** *In vitro* model and characterization of cellular and single-EV PD-L1 protein. **a** Immunofluorescence staining of PD-L1 protein in H1568 cells with/without interferon-gamma (IFN-γ) stimulation. Cell nuclei and PD-L1 protein were stained blue (by DAPI) and magenta (by anti-PD-L1 antibody), respectively. **b** Original TIRF microscopic images for Fig. 2c. The insets show the PD-L1 protein signals on a single EV. **c** A comparison of two different anti-PD-L1 antibodies provided by Cell Signaling Technology and Abcam to detect PD-L1 proteins on the surface of single EVs derived from IFN-γ-stimulated H1568 cells with $^{Au}$SERP. The H1568 EVs were spiked in healthy donor EVs at a 1:1 ratio with $10^9$ particles/mL each. The data were expressed as mean ± SD; n = 2. TFI, total fluorescence intensity; a.u., arbitrary units. **d** Representative TIRF microscopic images of PD-L1 protein expression of the H1568 single EVs stained by the two different antibodies. The insets show the PD-L1 protein signals on a single EV.

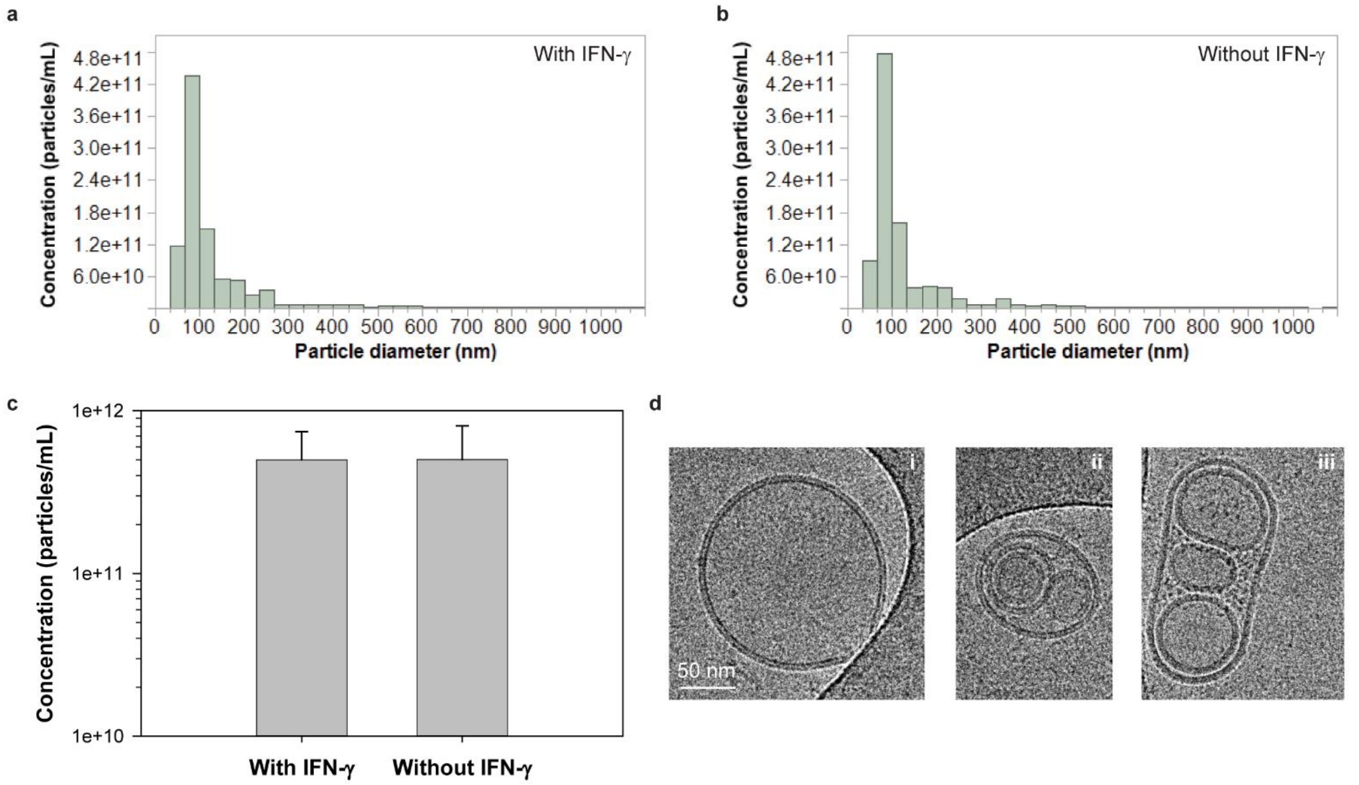

**Fig. S3.** Size and morphological characterization of EVs produced by H1568 cells. **a-b** Size distributions of EVs derived from H1568 cells with/without IFN-γ stimulation measured with Tunable Resistive Pulse Sensing (TRPS). **c** A comparison of their EV concentrations. The data were expressed as mean ± SD; n = 3. **d** Cryo-TEM images of EVs derived from IFN-γ-stimulated H1568 cells.

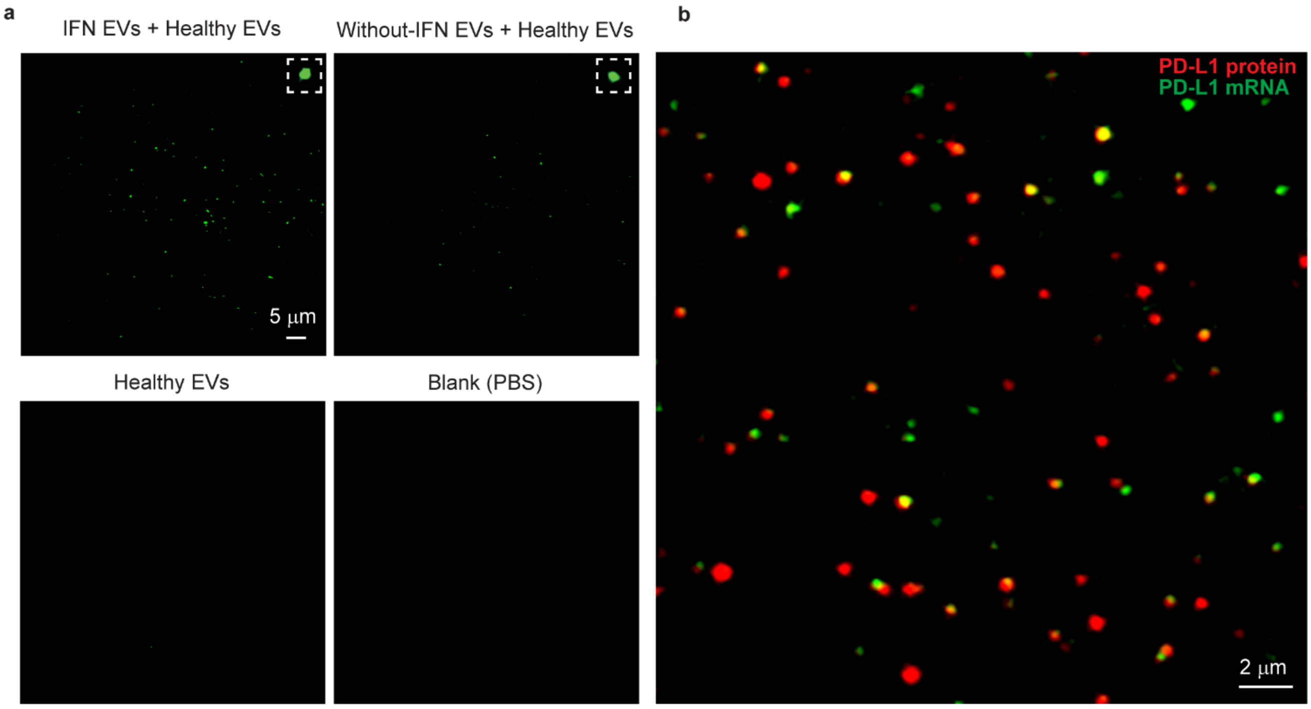

**Fig. S4.** *In vitro* model and characterization of single-EV PD-L1 protein and mRNA with $^{Au}$SERP. **a** Original TIRF microscopic images for Fig. 3e. The insets show the PD-L1 mRNA signals on a single EV. **b** Multiplex imaging of PD-L1 protein and mRNA from single EVs derived from IFN-γ-stimulated H1568 cells. PD-L1 protein and mRNA were stained green and red, respectively. The colocalized single-EV PD-L1 protein and mRNA signals appear yellow.

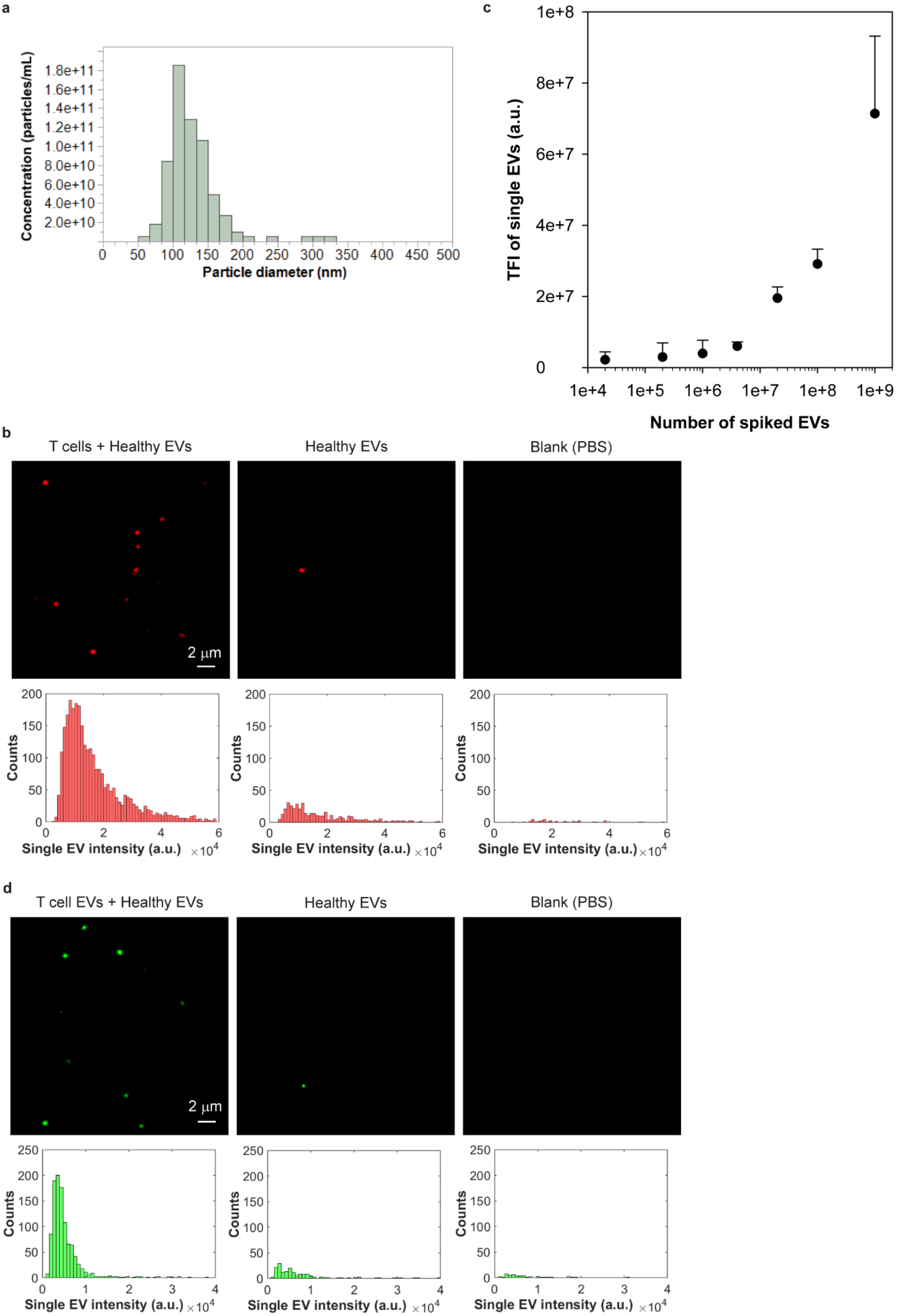
a
Concentration (particles/mL)
1.8e+11
1.6e+11
1.4e+11
1.2e+11
1.0e+11
8.0e+10
6.0e+10
4.0e+10
2.0e+10
0 50 100 150 200 250 300 350 400 450 500
Particle diameter (nm)
c
TFI of single EVs (a.u.)
1e+8
8e+7
6e+7
4e+7
2e+7
0
1e+4 1e+5 1e+6 1e+7 1e+8 1e+9
Number of spiked EVs
b
T cells + Healthy EVs
Healthy EVs
Blank (PBS)
2 μm
Counts
Single EV intensity (a.u.)
×10^4
d
T cell EVs + Healthy EVs
Healthy EVs
Blank (PBS)
2 μm
Counts
Single EV intensity (a.u.)
×10^4

**Fig. S5.** *In vitro* model and characterization of single-EV PD-1 protein and mRNA. **a** The size distribution of EVs produced by activated T cells measured with TRPS. **b** Representative TIRF microscopic images and their corresponding histograms of PD-1 protein expression on the surface of single EVs derived from activated T cells in comparison with negative controls. The single-EV PD-L1 protein signals were characterized with $^{Au}$SERP using an anti-PD-1 antibody and the TSA method. T cell-derived EVs were spiked in healthy donor EVs with 5 × $10^{10}$ particles/mL each. TFI, total fluorescence intensity; a.u., arbitrary units. **c** Quantitative detection of PD-1 protein with $^{Au}$SERP. T cell-derived EVs were spiked in healthy donor EVs at different concentrations ranging from 0 to 5 × $10^{10}$ particles/mL, while the healthy donor EV concentration was kept constant at 5 × $10^{10}$ EVs/mL. The limit of detection (LOD) of $^{Au}$SERP for PD-1 protein was ~ $10^6$ spiked T cell EVs. The data were expressed as mean ± SD; n = 3. **d** Representative TIRF microscopic images and their corresponding histograms of PD-1 mRNA in single EVs derived from activated T cells in comparison with negative controls. The single-EV PD-L1 mRNA signals were characterized with $^{Au}$SERP using PD-1 CLN-MBs. T cell-derived EVs were spiked in healthy donor EVs with 5 × $10^{10}$ particles/mL each.

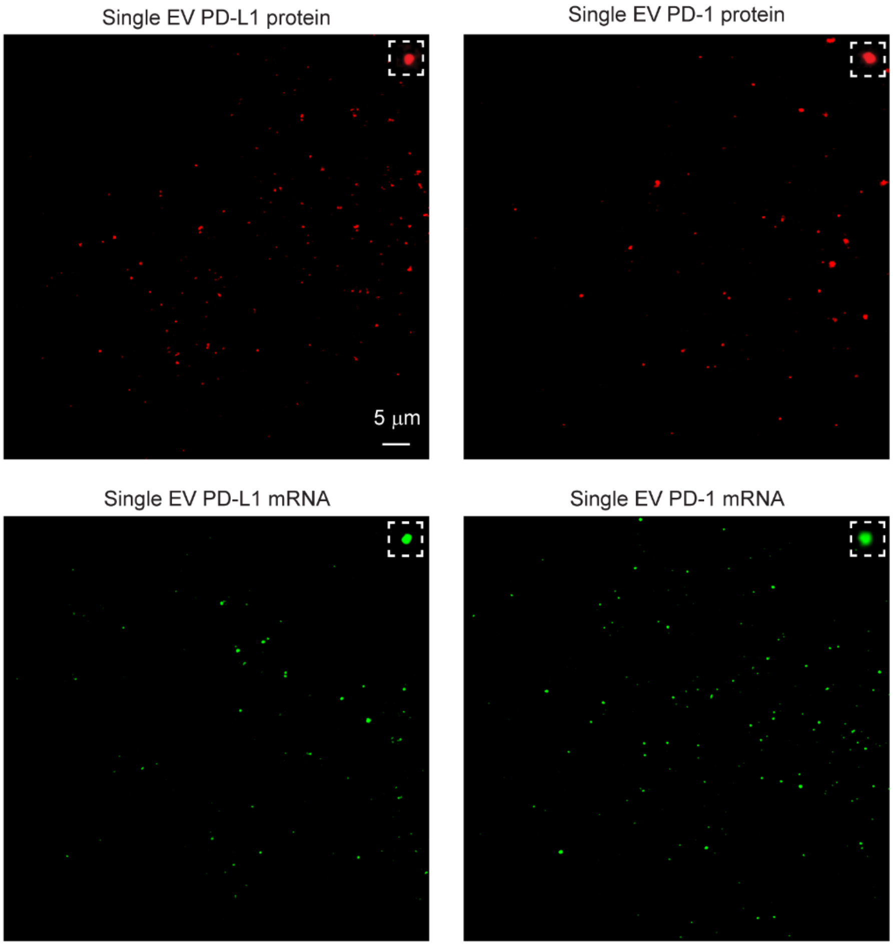

**Fig. S6.** Original TIRF microscopic images for Fig. 4c. The insets show the PD-1/PD-L1 protein and mRNA signals on a single EV.

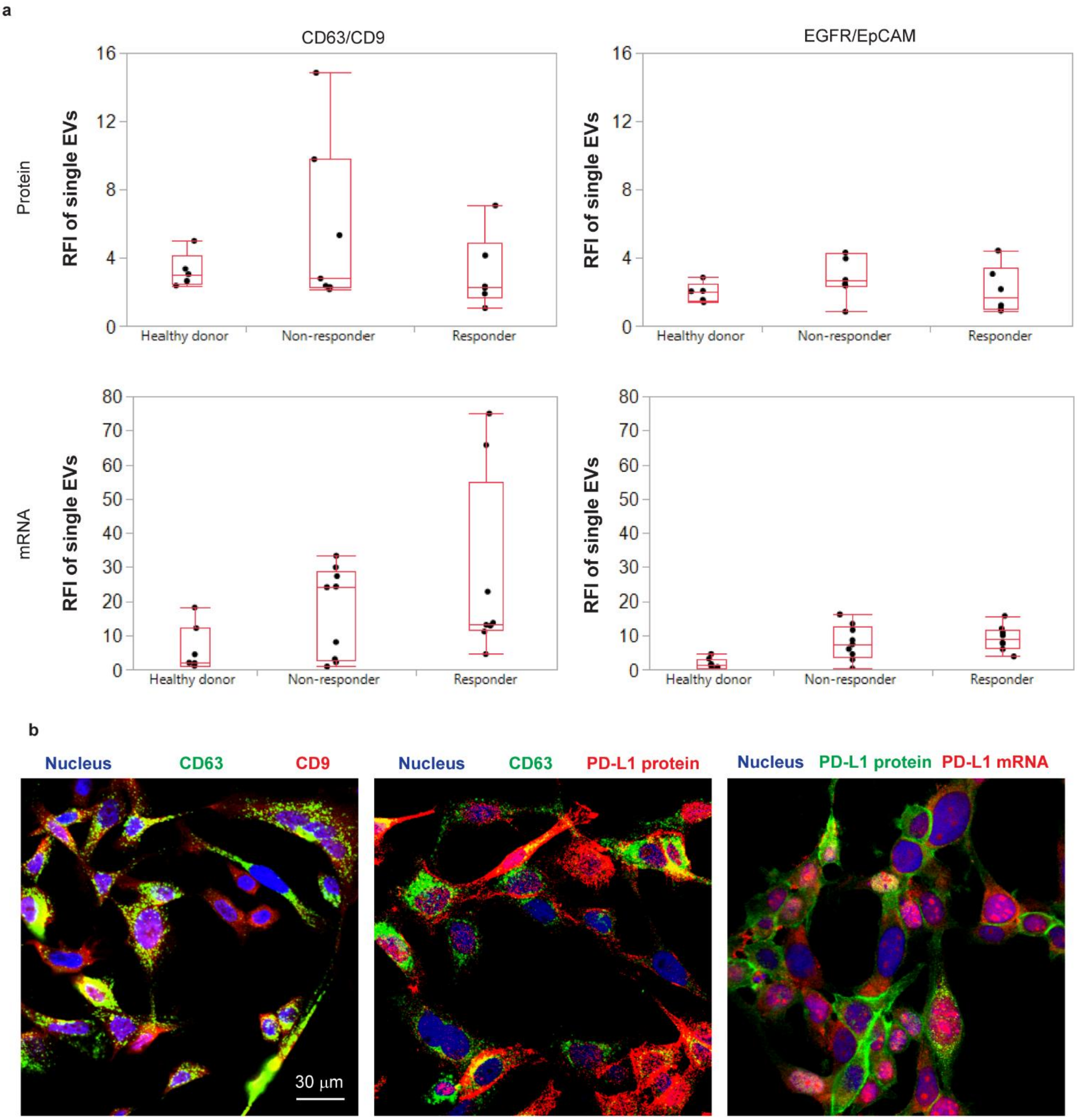

**Fig. S7.** Single-EV PD-L1 protein and mRNA characterization in subpopulations. **a** Comparisons of two different capture antibody cocktails (anti-CD63/CD9 and anti-EGFR/EpCAM) on the measurements of PD-L1 protein and mRNA signals in single EVs isolated from the serum of healthy donors, non-responders, and responders with $^{Au}$SERP. Each antibody (anti-CD63, anti-CD9, anti-EGFR, and anti-EpCAM) was used at a concentration of 10 μg/mL. For protein characterization, a cohort including healthy donors (n = 5), non-responders (n = 7), and responders (n = 6) was tested. For mRNA characterization, a cohort including healthy donors (n = 7), non-responders (n = 9), and responders (n = 8) was tested. RFI, relative fluorescence intensity.

**b** Confocal fluorescence microscopic images of CD63/CD9/PD-L1 proteins and PD-L1 mRNA in IFN-γ-stimulated H1568 cells. CD63/CD9/PD-L1 proteins were stained using the corresponding antibodies, while PD-L1 mRNAs were visualized using PD-L1 MBs via fluorescent *in situ* hybridization. Cell nuclei were stained blue using DAPI.

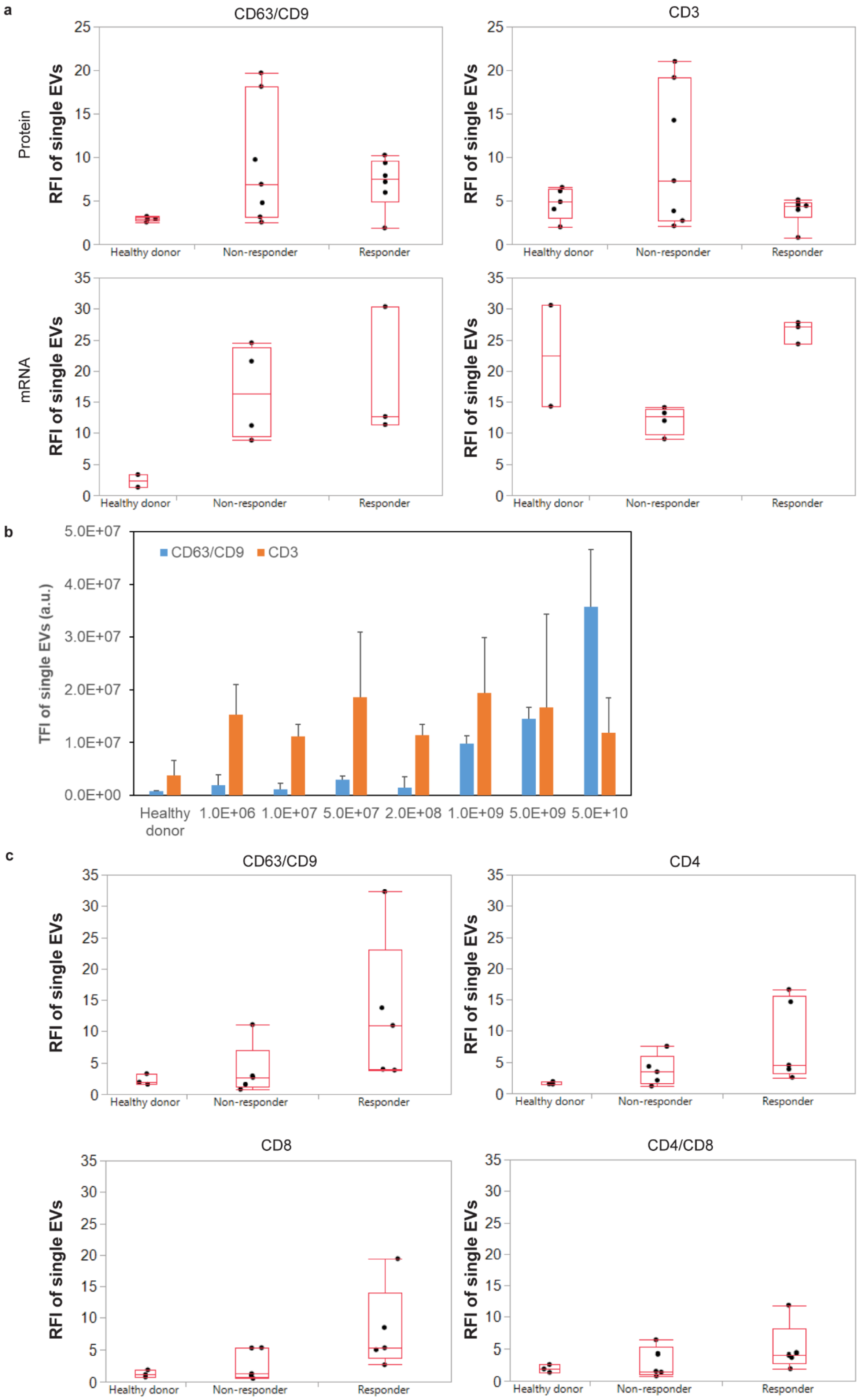
a
CD63/CD9
CD3
Protein
mRNA
RFI of single EVs
Healthy donor
Non-responder
Responder
0
5
10
15
20
25
30
35
b
TFI of single EVs (a.u.)
CD63/CD9
CD3
0.0E+00
1.0E+07
2.0E+07
3.0E+07
4.0E+07
5.0E+07
Healthy donor
1.0E+06
1.0E+07
5.0E+07
2.0E+08
1.0E+09
5.0E+09
5.0E+10
c
CD63/CD9
CD4
CD8
CD4/CD8

**Fig. S8.** A comparison of different capture antibodies on the measurement of PD-1 protein and mRNA signals in single EVs with $^{Au}$SERP. **a** A comparison of anti-CD63/CD9 and anti-CD3 capture antibodies on the measurement of PD-1 protein and mRNA signals in single EVs isolated from sera of healthy donors, non-responders, and responders. The cocktail of anti-CD63/CD9 antibodies was used at a concentration of 10 µg/mL each, while the anti-CD3 antibody was used at 20 µg/mL. For protein characterization, a cohort including healthy donors (n = 5), non-responders (n = 7), and responders (n = 6) was tested. For mRNA characterization, a cohort including healthy donors (n = 2), non-responders (n = 4), and responders (n = 3) was tested. **b** Quantitative detection of PD-1 protein with different capture antibodies (anti-CD63/CD9 and anti-CD3). T cell-derived EVs were spiked in healthy donor EVs at different concentrations ranging from 0 to $5 \times 10^{10}$ particles/mL, while the healthy donor EV concentration was kept constant at $5 \times 10^{10}$ EVs/mL. TFI, total fluorescence intensity; a.u., arbitrary units. **c** A comparison of different capture antibodies (anti-CD63/CD9, anti-CD4, anti-CD8 and anti-CD4/CD8) on the measurement of PD-1 protein on the surface of single EVs isolated from the serum of healthy donors (n = 3), non-responders (n = 5), and responders (n = 5). The cocktail of anti-CD63/CD9 and anti-CD4/CD8 antibodies were both used at a concentration of 10 µg/mL for each antibody, while single anti-CD4 and anti-CD8 antibodies were used at 20 µg/mL.

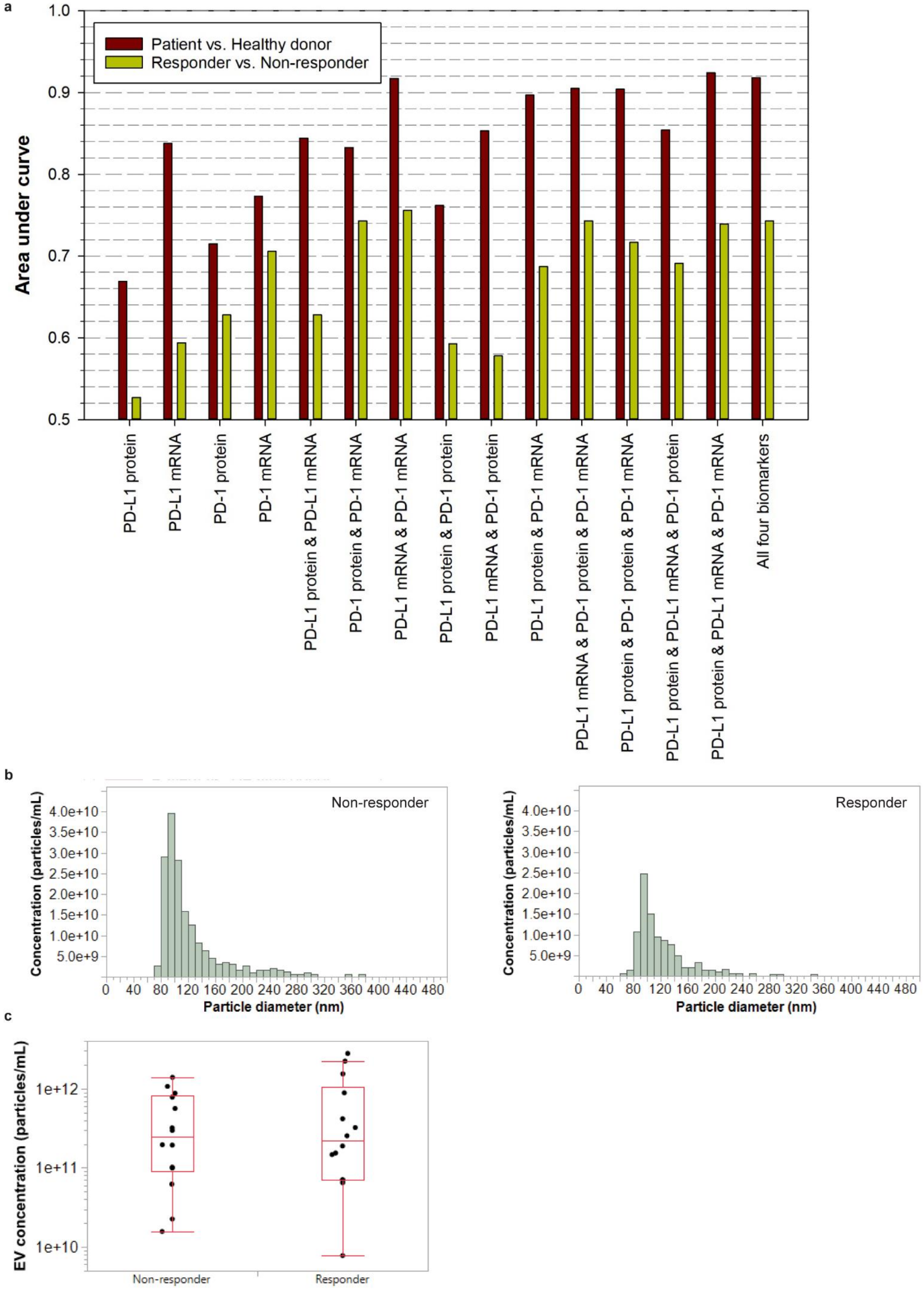
a
Area under curve
1.0
0.9
0.8
0.7
0.6
0.5
Patient vs. Healthy donor
Responder vs. Non-responder
PD-L1 protein
PD-L1 mRNA
PD-1 protein
PD-1 mRNA
PD-L1 protein & PD-L1 mRNA
PD-1 protein & PD-1 mRNA
PD-L1 mRNA & PD-1 mRNA
PD-L1 protein & PD-1 protein
PD-L1 mRNA & PD-1 protein
PD-L1 protein & PD-1 mRNA
PD-L1 mRNA & PD-1 protein & PD-1 mRNA
PD-L1 protein & PD-1 protein & PD-1 mRNA
PD-L1 protein & PD-L1 mRNA & PD-1 protein
PD-L1 protein & PD-L1 mRNA & PD-1 mRNA
All four biomarkers
b
Non-responder
Responder
Concentration (particles/mL)
4.0e+10
3.5e+10
3.0e+10
2.5e+10
2.0e+10
1.5e+10
1.0e+10
5.0e+9
0 40 80 120 160 200 240 280 320 360 400 440 480
Particle diameter (nm)
c
EV concentration (particles/mL)
1e+12
1e+11
1e+10
Non-responder
Responder

**Fig. S9.** Characterization of EVs from the serum of non-responders (n = 27) and responders (n = 27). **a** Area under the curve (AUC) values of receiver operating characteristic (ROC) curve analyses for the single-EV immunotherapy biomarkers measured with $^{Au}$SERP. For NSCLC diagnosis, the sample size consisted of patients (n = 54) vs. healthy donors (n = 20). For prediction of NSCLC patient response to anti-PD-1/PD-L1 immunotherapy, the sample size consisted of responders (n = 27) vs. non-responders (n = 27). An ROC curve analysis was performed for a single biomarker and multiple biomarkers with different permutations. **b** Representative size distributions of the patient serum-derived EVs measured with TRPS. **c** Box plots of the concentrations of the patient serum-derived EVs measured with TRPS.

**Table S1.** Clinical characteristics of stage IV NSCLC patients.

| | **Non-responder (n= 27)** | **Responder (n = 27)** |
|---|---|---|
| Age | | |
| Median (range) | 63 (47 – 83) | 65 (45 – 86) |
| Gender | | |
| Male | 14 | 18 |
| Female | 13 | 9 |
| Race | | |
| Caucasian | 23 | 24 |
| African American | 4 | 3 |
| Smoking history | | |
| Current | 7 | 6 |
| Former | 19 | 21 |
| Never | 1 | 0 |
| Performance status | | |
| 0 | 5 | 6 |
| 1 | 18 | 19 |
| 2 | 4 | 2 |
| Histology | | |
| Adenocarcinoma | 19 | 18 |
| Squamous cell | 7 | 5 |
| Adenosquamous | 0 | 1 |
| NOS | 1 | 3 |
| PD-L1 IHC | | |
| Positive | 12 | 17 |
| Negative | 6 | 4 |
| Unknown | 9 | 6 |
| EGFR | | |
| Wild | 18 | 22 |
| Mutant | 5 | 1 |
| Unknown | 4 | 4 |
| Drug | | |
| Nivolumab | 20 | 17 |
| Pembrolizumab | 7 | 9 |
| Atezolizumab | 0 | 1 |

**Table S2.** Antibodies used for single-EV capture and detection.

| **Antibody** | | **Catalog number/ Brand name** | **Supplier** |
|---|---|---|---|
| Capture | CD63 | MAB5048 | R&D Systems |
| | CD9 | MAB1880 | R&D Systems |

| | | | |
|---|---|---|---|
| | EGFR (Cetuximab) | Erbitux® | ImClone LLC |
| | EpCAM | AF960 | R&D Systems |
| | CD3 | MAB100100 | R&D Systems |
| | Biotinylated CD4 | 344610 | BioLegend |
| | Biotinylated CD8 | 344720 | BioLegend |
| Detection | CD63 - Alexa Fluor® 488 | sc-5275 AF488 | Santa Cruz Biotechnology |
| | PD-L1 | 86744S | Cell Signaling Technology |
| | PD-L1 | ab205921 | Abcam |
| | PD-1 | 86163S | Cell Signaling Technology |
